# Comparison of two African rice species through a new pan-genomic approach on massive data

**DOI:** 10.1101/245431

**Authors:** Cécile Monat, Christine Tranchant-Dubreuil, Stefan Engelen, Karine Labadie, Emmanuel Paradis, Ndomassi Tando, François Sabot

## Abstract

Pangenome theory implies that individuals from a given group/species share only a given part of their genome (core-genome), the remaining part being the dispensable one. Domestication process implies a small number of founder individuals, and thus a large core-genome compared to dispensable at the first steps of domestication. We sequenced at high depth 120 cultivated African rice *Oryza glaberrima* and of 74 wild relatives *O. barthii*, and mapped them on the external reference from Asian rice *O. sativa*. We then use a novel DepthOfCoverage approach to identif missing genes. After comparing the two species, we shown that the cultivated species has a smaller core-genome than the wild one, as well as an expected smaller dispensable one. This unexpected output however replaces in perspective the inadequacy of cultivated crops to wilderness.

## Introduction

Pan-genome [Tettelin et al., 2005] is defined as the whole set of DNA sequence present in a given group such as a species. This pan-genome is further divided in three compartments:(*i*) the core-genome, composed of sequences present in all individuals of the group; (*ii*) the dispensable/variable/accessory-genome, sequences present in some but not all of the individuals; and (*iii*) the individual-specific genome, accounting for sequences present only in a single individual of the group. It has been first defined on microorganism [Tettelin et al., 2005] and is now commonly used on bacteria with some specifics tools developed for this type of analysis [Laing et al., 2010, Snipen and Ussery, 2010, Zhao et al., 2012, Sarovich and Price, 2014, Nieselt, 2015, Thakur and Guttman, 2016]. Few studies also started on other genera like animals [Boussaha et al., 2015], Human [Li et al., 2010], viruses [Aherfi et al., 2014], algae [Wang et al., 2014], fungi [Upadhyaya et al., 2015] and plants [Cheung et al., 2009, ?, Montenegro et al., 2017].

From this comes the idea that each individual has differences not only in term of alleles but also in term of sequence numbers and composition [Chia et al., 2012, Nieselt, 2015,Lu et al., 2015]. Moreover two genomes are similar not only by the sequences which they share, but also because they miss the same sequences [Snipen and Ussery, 2010]. These type of studies also confirm that having a single reference genome per species is certainly helping but is not enough to represent the whole genomic diversity of a species, as the reference genome has sequences absent in other individual of the same species and miss some other ones [Weigel and Mott, 2009, Gan et al., 2011, Upadhyaya et al., 2015].

Pan-genomic is thus a new way to understand evolution processes (such as sympatric speciation), and to establish new link between individuals (for instance, the link between two sub-species) as well as between different species.

One interesting aspect is to better characterize and understand the domestication process and its effects on genome, especially on genome diversity and structure.

Several studies showed that the analysis of several genomes belonging to the same species allowed to discover more intra-species diversity than expected [Tettelin et al., 2008]. Therefore, doing comparative genomics of pan-genomic type put forward the interest to work within the same species to represent a diversity which would not be necessarily visible by looking only at the inter-species level [Lukjancenko et al., 2012].

Rice is the most cultivated cereal for human consumption, with two species cultivated, the Asian rice *Oryza sativa*, worldwide used, and the African rice *Oryza glaberrima*, endemic to West Africa. The latter and its wild relative *Oryza barthii* are known to have a very low diversity (SNPs based) [Nabholz et al., 2014] compared to other grasses. This characteristic makes them good models to study the genomic diversity in light of pan-genome as with a reasonable number of individuals will cover the entire diversity of the species. For that purpose we resequenced 194 accessions of African rices (120 of the cultivated species and 74 of its wild relative) with Illumina technology and shown huge genic variations during domestication.

## Materials and Methods

### Plant material

One hundred and twenty accessions of *Oryza glaberrima* and 74 accessions of *Oryza barthii* were used in the study (see annexe E) and described in [Cub, 2017]. Plants were grown at IRD greenhouses at Montpellier (France) under normal conditions, and DNA extracted using *QIAGEN* DNAeasy kit (Germany) as recommended by suppliers. Accessions were chosen to span the geographical distribution as well as the phylogenetic tree defined with specific *Oryza glaberrima* markers [Orjuela et al., 2014] (Figs. S1, S2 and S3).

### DNA Sequencing

DNA was sequenced at the Genoscope (Evry, France) on Illumina HiSeq 2000/2500/400 with paired-end reads, 100 bases long, and cleaned as described in [Cub, 2017,?]. Briefly, the mean sequencing depth is of 35x with a maximum of 52x and a minimum of 21x (for more details see Table S1), representing a total of 7,749,300,142 and 4,378,115,069 reads for *O. glaberrima* and *O. barthii*, respectively (for more details see Table S2).

### Short reads mapping

Cleaned reads were then mapped against the Asian reference genome (Os-Nipponbare-Reference-IRGSP-1.0/MSU7.0 [Mcnally et al., 2009, Kawahara et al., 2013]) through a TOGGLE pipeline [Monat et al., 2015, ?], as follows: FASTQC control, repairing step to mate the pairs, mapping with BWA aln/sampe legacy [Li and Durbin, 2009] (edit distance of 5 bases), conversion into BAM file with Picardtools (!!REF!!), extraction of properly and unproperly mapped in pair data with SAMtools [Li et al., 2009]. Properly mapped in pair reads were subject to local realignment using GATK [Mckenna et al., 2010], duplicates removed with Picardtools (for details information about mapping see Percentage of mapping versus *Oryza sativa* reference genome for the whole accessions).

### Reads count and normalization

The reads count for each individual on different data sources (annotation line from the Asian rice (http://rice.plantbiology.msu.edu/)) were generated with the multicov tool from BEDTOOLS [Quinlan, 2014]. Counts were normalized to 1 million total reads according to the initial individual sequencing coverage and to a 1 kb bin length to obtain FPKM values (Features per kilobase of ‘genes’ per millions of reads).

### Pnorm and FDR analysis

Pnorm analysis and FDR approach at 5% were performed using R and specific packages pnorm and qvalue [with contributions from Andrew J. Bass et al., 2015]. Annotations or bins for which the normalized reads count mean were lower than 2 were arbitrarily previously ride out the analysis and considered as absent for all individuals. Pan-matrix obtained shown ‘0’ when gene/bin was considered as absent, ‘1’ if the gene/bin was considered as present and ‘Uk’ (Unknown) if feature does not pass the initial pnorm test.

### GO enrichment analyses

Gene ontology enrichment analyses were performed using a home-made R script and the topGO package [Alexa and Rahnenfuhrer, 2016], with a display of the 5 top most significant GO retrieved using the Fisher classic and weight01 algorithms.

### Identification of domesticated related gene families

Data from gene families were obtained from Green Phyl V.4 [Conte et al., 2008a, Conte et al., 2008b, Rouard et al., 2011], selecting sequences without spliced forms.

### Availability of scripts

The whole Perl and R scripts to perform:

- normalization;
- chromosomal, individual and population effects checking;
- Pnorm and FDR analysis;
- assignation of compartment;
- GO and gene families analysis;

are available on github under Cecill-B/GPLv3 double licenses.

## Results

### Mapping and Normalization

We sequenced 163 accessions of the cultivated African rice *O. glaberrima* and 86 accessions of its wild relative *O. barthii* using high depth Illumina technology. After cleaning the raws reads were mapped against the Asian reference genome (Os-Nipponbare-Reference-IRGSP-1.0 [Mcnally et al., 2009, Kawahara et al., 2013]) as an external reference to avoid biases to any of the two species. The mean percentage of mapping for *O. glaberrima* is 86.49% (81.57% for properly paired reads) and 86.44% (81.45%) for *O barthii*.

We then performed reads count normalization according to the initial reads number specific to each individual reported to 1 million and the gene/bin length reported to 1kb, obtaining numerical FKPM data. Distribution of FPKM density mainly follows a normal law (see Fig. 1), thus we applied successively pnorm and FDR statisticals test to define if a given gene/bin was present or not for each individual.

**Figure 1.**
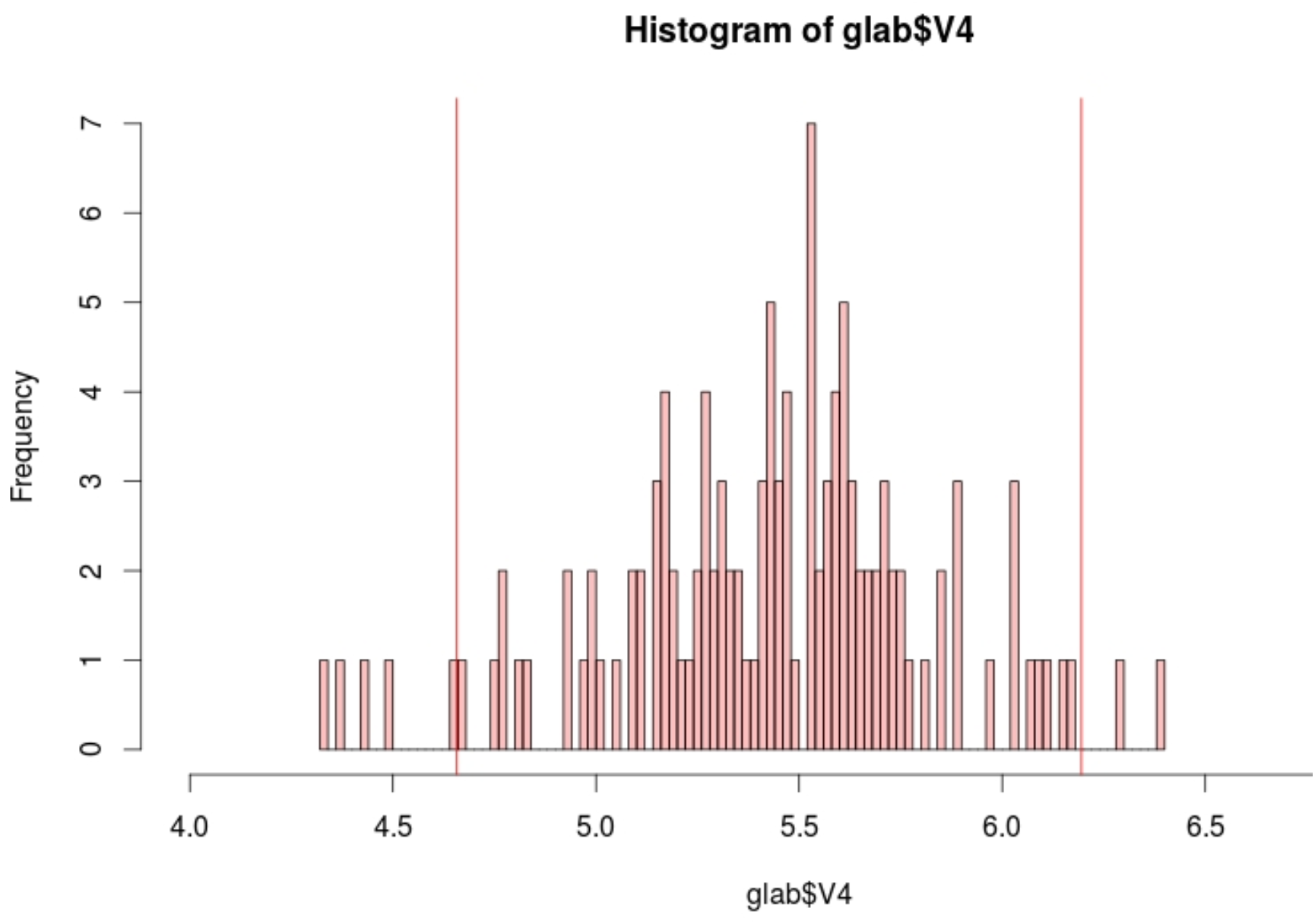
Reads distribution for one feature on *O. glaberrima*

### Chromosomal, individual and population effects

First, using reference sequence annotation data, we checked whether the global profiles were the same between the different chromosomes, no matter if we look to gene, CDS, UTR or TE features and if there were or not individual(s) effects. The first test was to see if the profile were different according to the chromosome we looked at. To obtain those profil we looked for the outliers at differents thresholds. We seen that there are no significant differences between the 12 chromosomes neither on *O. glaberrima* nor on *O. barthii* (see Figs. S4 and S5). We then performed the same analysis only on features representing exons (see Figs. S6 and S7) and representing not exons (see Figs. S8 and S9) and confirm profils are the same no matter the type of annotation features we are looking at. As a proxy for the rest of the chromosome we choose the chromosome 10 for the following verifications (central in the little range of chromosomes profiles).

In order to test if there were individual effect, we performed a hundred bootstrap-like analysis choosing 76 individuals. No individual effect were detectable on *O. glaberrima* (see Fig. S10) but in *O. barthii* we found two classes of profils depending on the individuals in the group (see Fig. S11). As described by Orjuela et al [Orjuela et al., 2014], this species can be divided in two populations respectively with 23 and 51 individuals distributed as in Fig. S12. To check if the two profiles were due to population split, we tested the hundred bootstrap analysis on *O. barthii* with only 23 individuals of the first population (see Fig. S13) and 51 of the second (see Fig. S14). Two profiles were still detectable on the population 1, so we checked if 23 individuals was enough to determine these profiles. We performed the same analysis, only with the population 2 individuals but with bootstrap of 23 individuals. This time the curves show more discrepancy suggesting that 23 is not a sufficient number of individuals to see a robust profile (see Fig. S15). To be sure that it is necessary to have a minimum number of individuals to have a strong signal, we performed verifications on *O. glaberrima*. Both the 23 and 51 verifications conserved the robust profil (see Figs. S16 and S17).

### 0.1 Pan-genome compartments

The pan-genome of *O. glaberrima* is comprised into the core-genome of *O. barthii* as it represents 39,106 and 39,730 genes respectively (see Fig. 2). On the other hand, the dispensable-genome of the cultivated species is bigger than wild one as its represents 5,299 and 745 genes respectively (see Fig. 2, Table 1 and Table 2). We identified 5,258 genes (more than 10% of the repertoire of wild genes) common to the core-genome of *O. barthii* and to the dispensable-genome of *O. glaberrima*. On the other hand we found only 23 genes common to the dispensable-genome of the wild species and the core-genome of the cultivated one. We were not able to detect any individual-specific genes.

**Figure 2.**
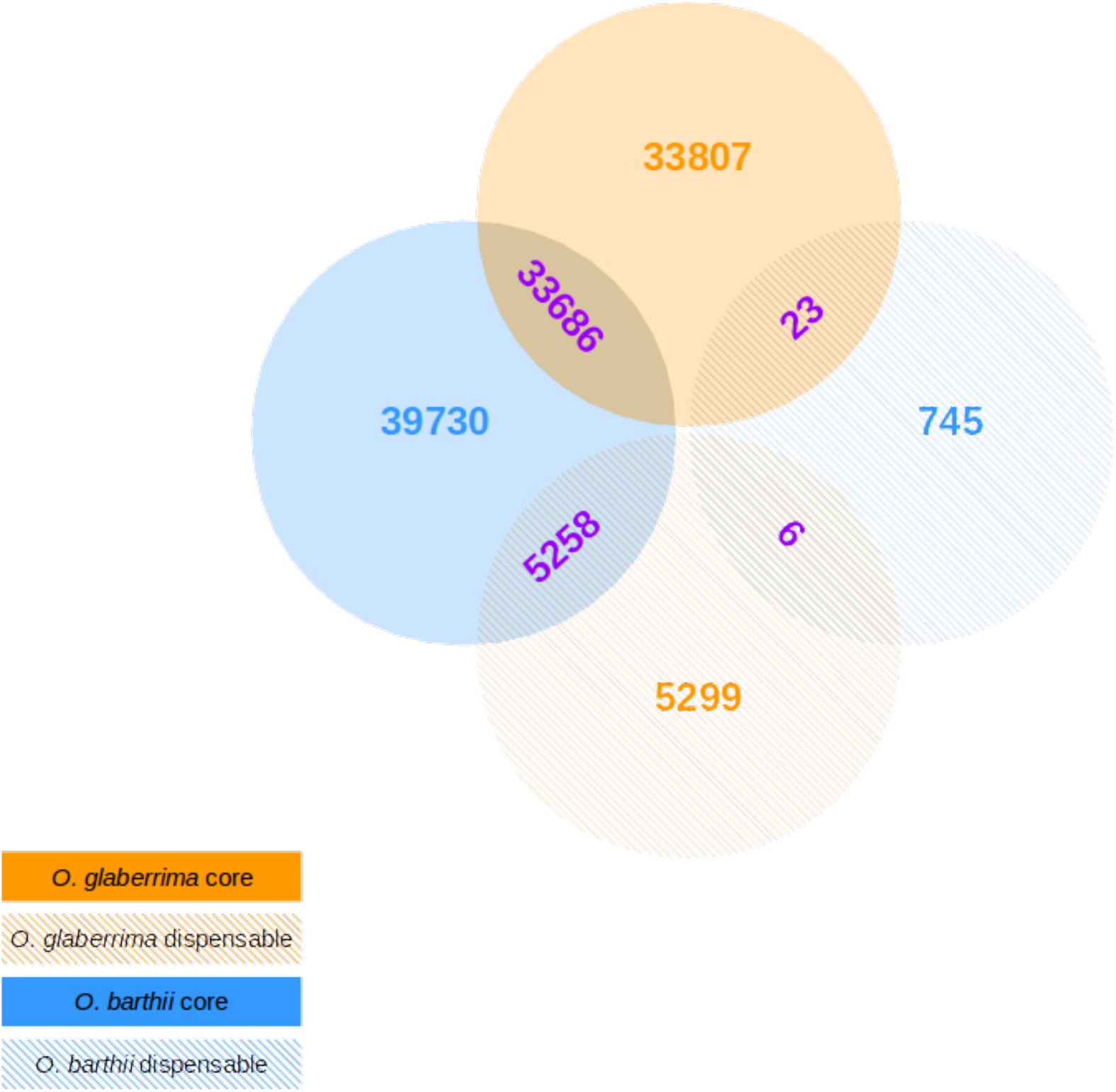
Pan-genome of *O. glaberrima* and *O. barthii*

**Table 1.**
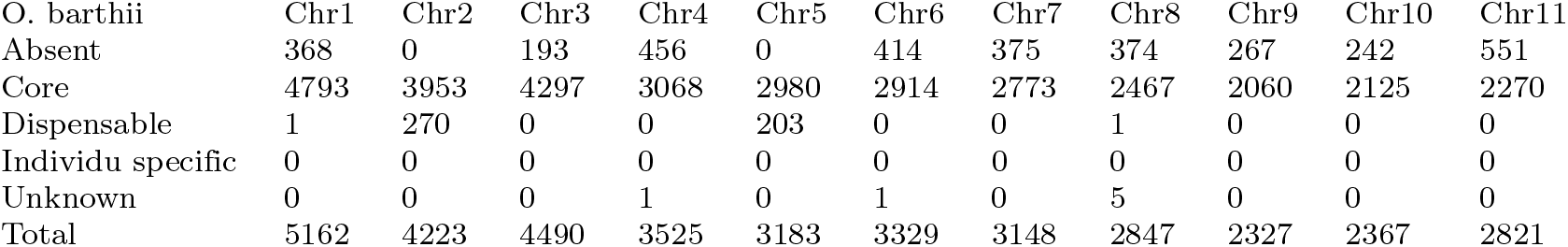
Pan-genome of *O. glaberrima*. The line first line represents the features we have taken apart because the reads count was not high enough to perform the analysis (mean ≤ 2). Absent mean is considered as zero for all the individuals. Core mean is considered as one for all the individuals. Dispensable mean is considered as zero for at least one individual. Individu specific mean is considered as one for only one individual. Unknown mean the p-value was not realisable on the reads count distribution for this particular feature.

**Table 2.**
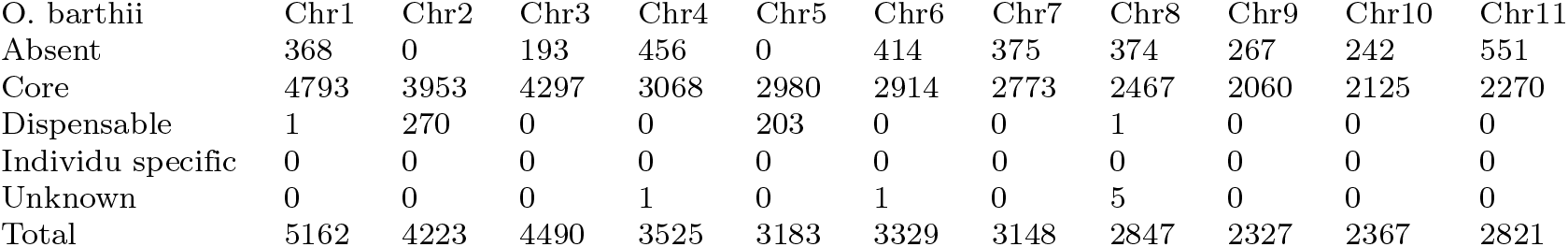
Pan-genome of *O. barthii*. Same legend as Table 1.

We looked to the distribution of genes compartment along the chromosomes as presented in Fig. 3. As expected the profile for the core and the dispensable genes are mirrored. The repartition of genes, no matter to which compartment they belong to, does not seem to follow a particular profile. It seems to be regular along the chromosomes.

**Figure 3.**
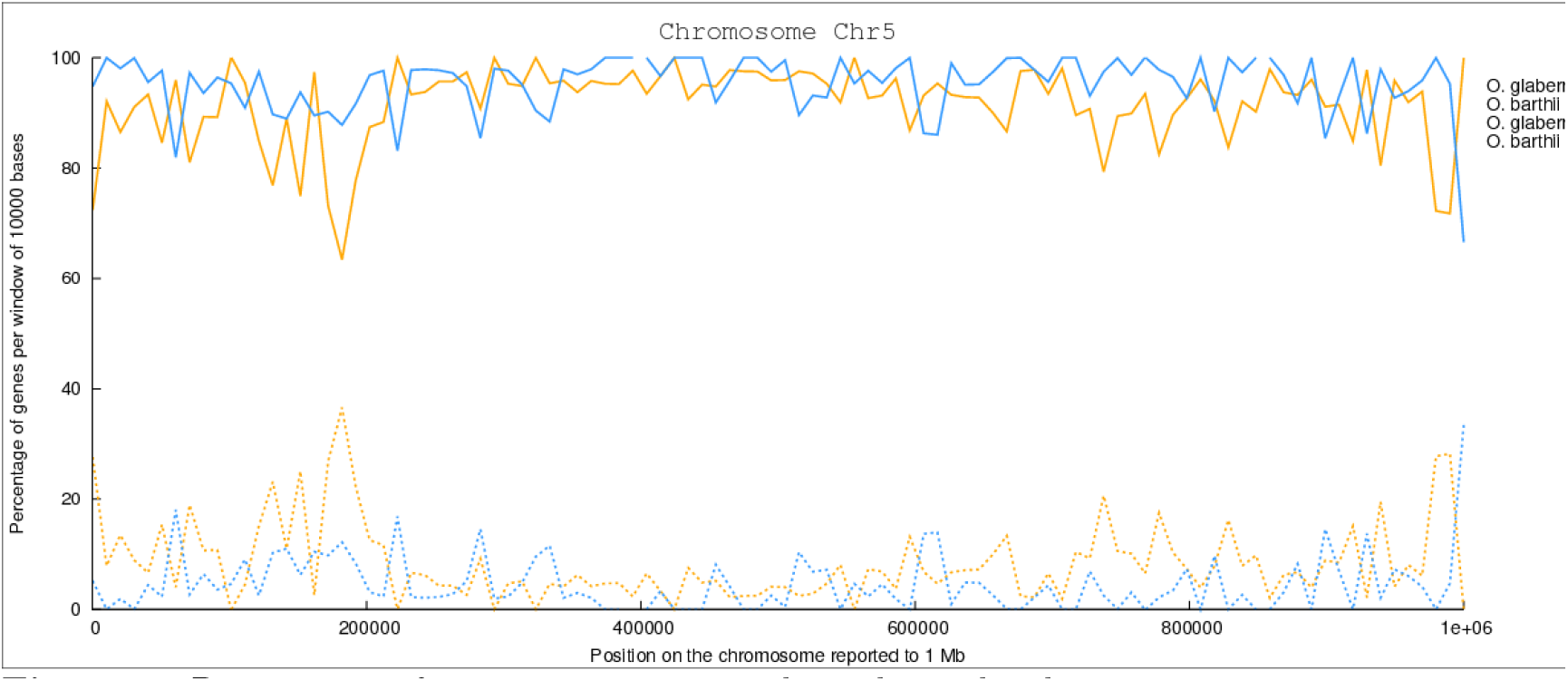
Repartition of gene compartments throughout the chromosome 5

### 0.2 Identification of domestication related ontology/ gene families

We found 10 GO changing of compartment between *O. barthii* and *O. glaberrima* (see Table 3). Six of them switch from the core-genome to the dispensable-genome, and four from the dispensable of *O. barthii* seem to be totally missing in *O. glaberrima*.

**Table 3.**
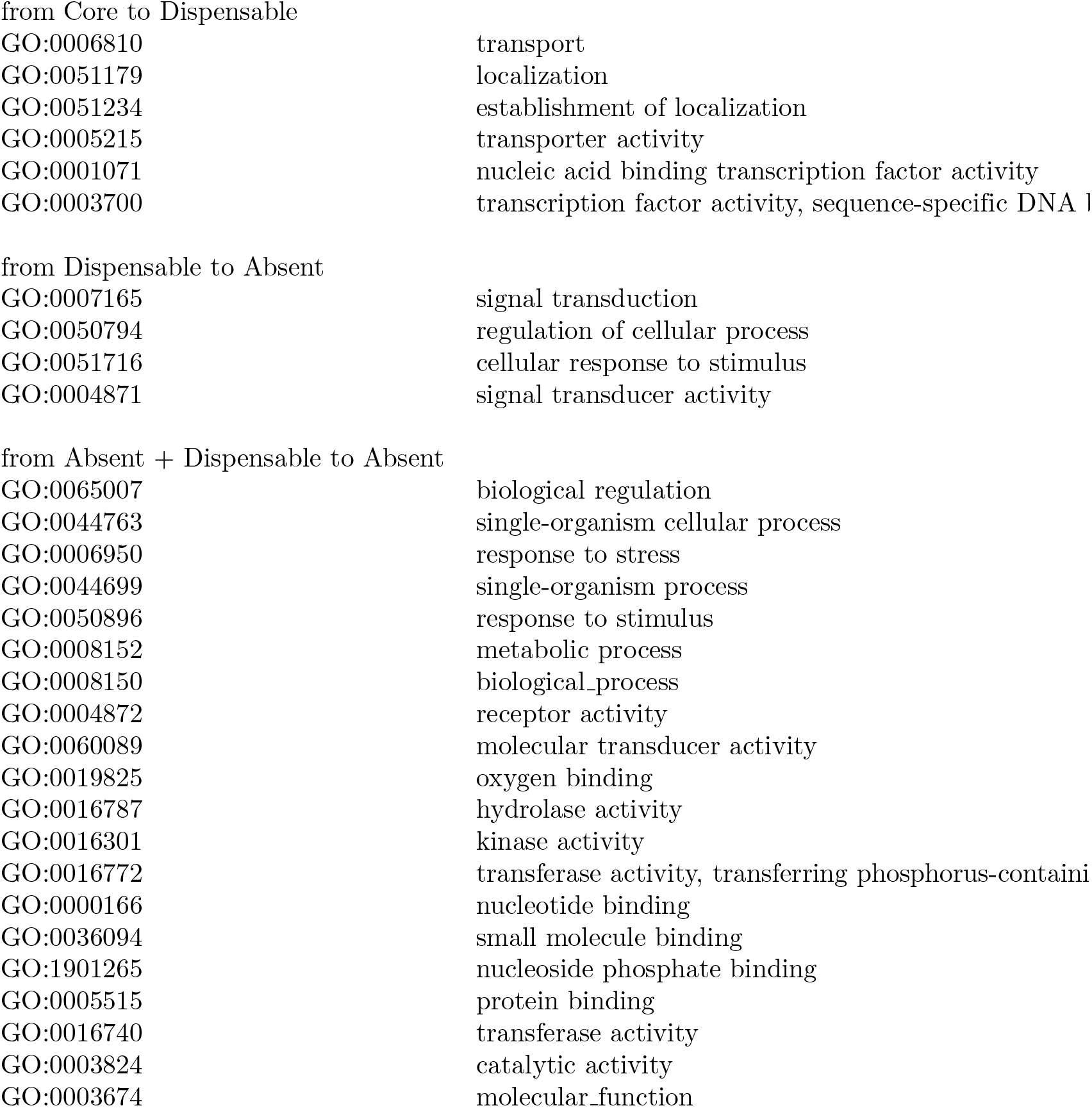
List of gene ontology movements from *O. barthii* to *O. glaberrima*

|  |  |
| --- | --- |
| GO:0006810 | transport |
| GO:0051179 | localization |
| GO:0051234 | establishment of localization |
| GO:0005215 | transporter activity |
| GO:0001071 | nucleic acid binding transcription factor activity |
| GO:0003700 | transcription factor activity, sequence-specific DNA binding |
| GO:0007165 | signal transduction |
| GO:0050794 | regulation of cellular process |
| GO:0051716 | cellular response to stimulus |
| GO:0004871 | signal transducer activity |

**Table 3.** List of gene ontology movements from *O. barthii* to *O. glaberrima*
|  |  |
| --- | --- |
| GO:0065007 | biological regulation |
| GO:0044763 | single-organism cellular process |
| GO:0006950 | response to stress |
| GO:0044699 | single-organism process |
| GO:0050896 | response to stimulus |
| GO:0008152 | metabolic process |
| GO:0008150 | biological process |
| GO:0004872 | receptor activity |
| GO:0060089 | molecular transducer activity |
| GO:0019825 | oxygen binding |
| GO:0016787 | hydrolase activity |
| GO:0016301 | kinase activity |
| GO:0016772 | transferase activity, transferring phosphorus-containing groups |
| GO:0000166 | nucleotide binding |
| GO:0036094 | small molecule binding |
| GO:1901265 | nucleoside phosphate binding |
| GO:0005515 | protein binding |
| GO:0016740 | transferase activity |
| GO:0003824 | catalytic activity |
| GO:0003674 | molecular function |

**Table 4.**
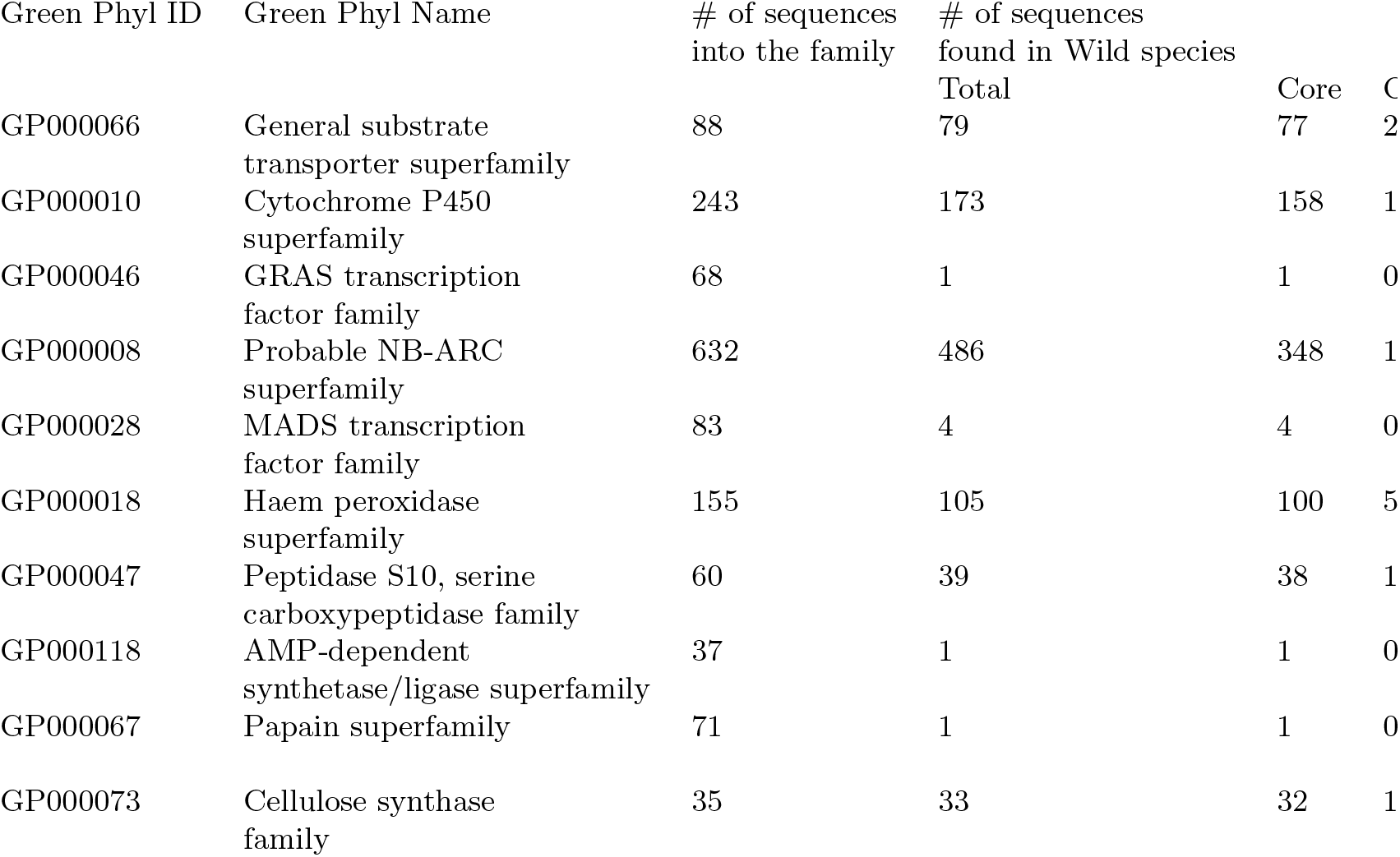
Gene families

**Table 5.**
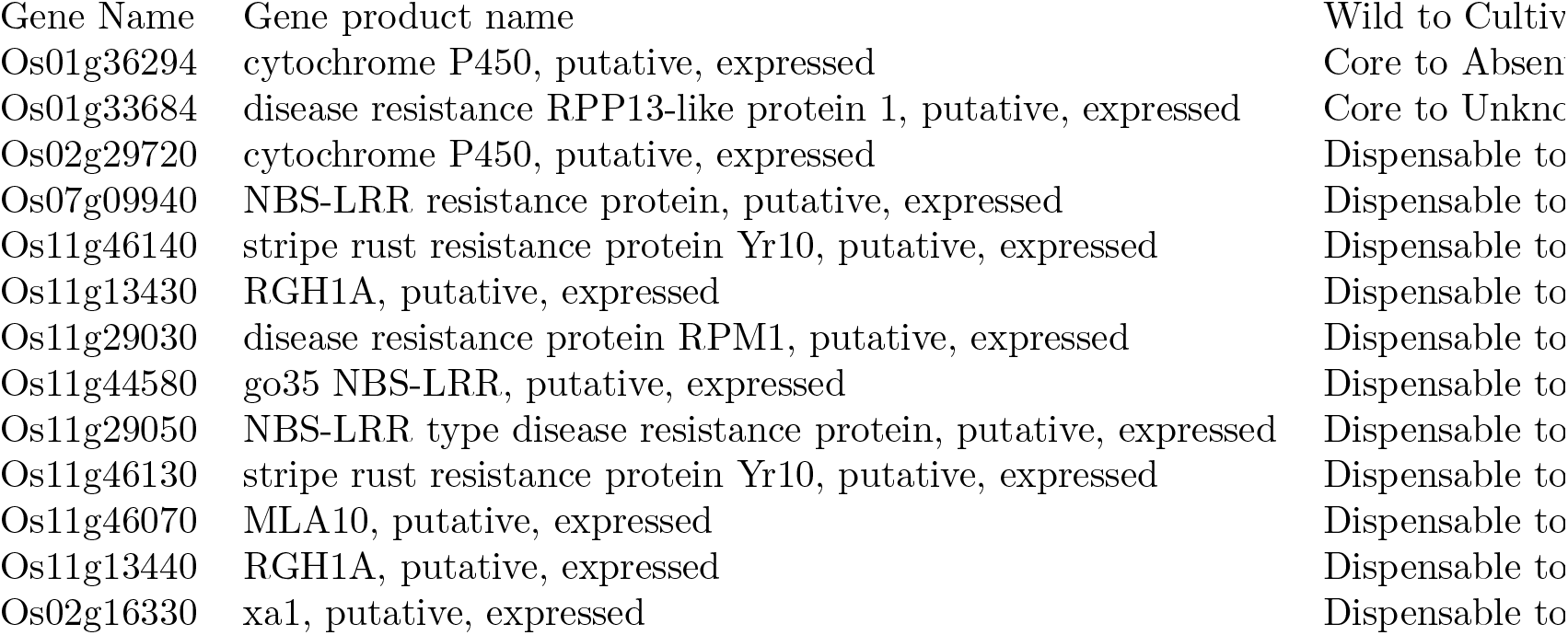
Gene families

**Table 6.**
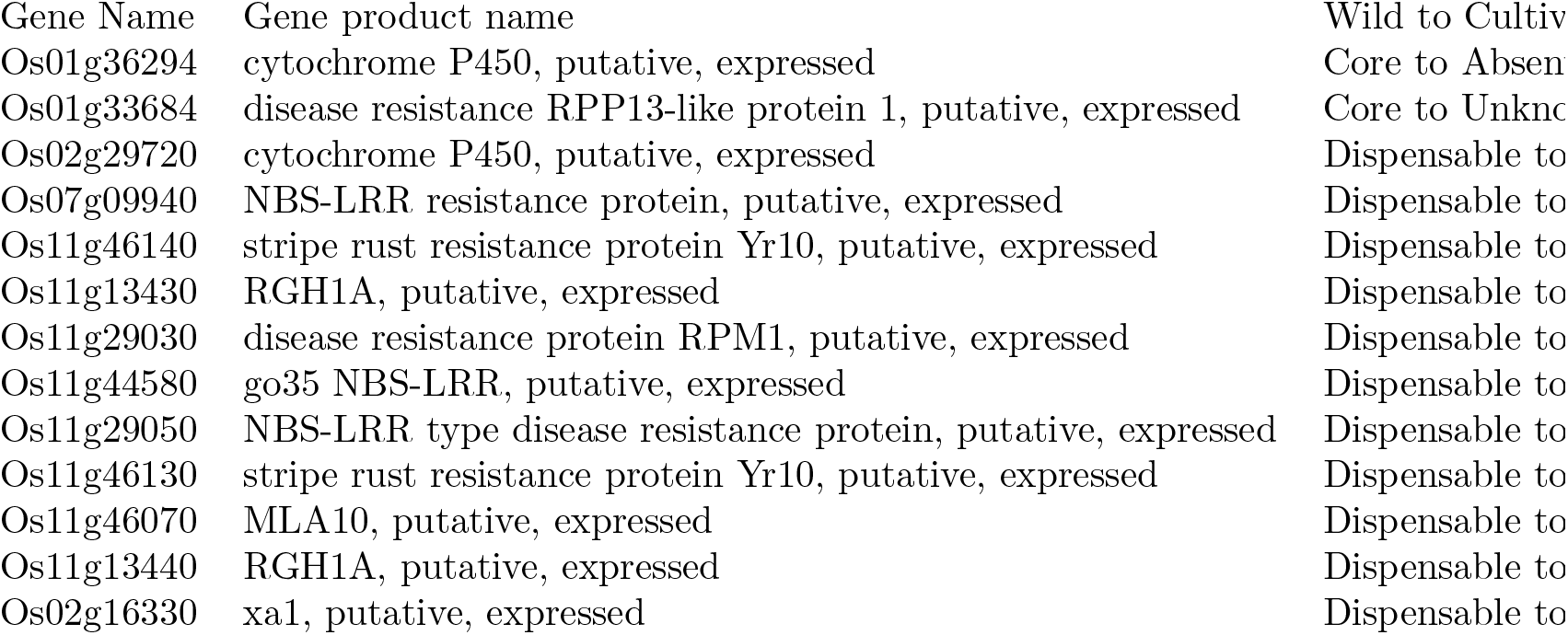
Genes belonging to gene family

We also found 20 GO for which the process of disappearing seems to start as those GO are totally missing in *O. glaberrima* but are present in both dispensable and absent part of *O. barthii* (see Fig. 3).

However as the informations given by the GO enrichment analysis was not clearly identifying something different between the two species, we decided to look at members of some gene families (Fig. 4). First of all we confirmed that the whole set of genes of the same family are not necessarily in the same compartment. Secondly we saw the same trend as we seen with GO, *i.e* from *O. barthii* core to *O. glaberrima* dispensable, from *O. barthii* dispensable to disapearing in *O. glaberrima* and from *O. barthii* missing or in dispensable-genome to complete disapearing in *O.glaberrima*. In addition we found also a new kind of movement (see Fig. 5) which are from *O. barthii* core to disapearing in *O. glaberrima* and from the *O. barthii* dispensable to the core-genome of *O.glaberrima* (see Fig. 6). One gene family give us an information about one ‘Uk’ genes as it was classified as ‘Uk’ in the cultivated but classified into the core-genome for the other one.

## Discussion

Pan-genomes studies allow to have a new point of view about diversity of a species as well as to have new insights on comparative genomic between two or more species. Pan-genome classes genes into three compartments according to the number of individuals which carry it in the sample studied. Core genes are often related to essential functions like development, dispensable genes to environmental adaptation and biotic stress. Those genes were presented to be really important into the diversification process. Domestication is known to reduce diversity from the wild to the cultivated species in term of SNPs. We are interested to see the effect of domestication process on the pan-genome structure. To answer this question we performed pan-genome analyses on both the African cultivated rice and its wild relative.

It is important to remember that in pan-genomic studies, results and conclusions are really dependent on many criteria. First of all, the results clearly depend on the sample and the individuals used. In this study we have resequenced accessions all along the diversity of the two species, trying also to have a good geographic representation of these accessions. We have tried to reduce to its minimal the bias that individual sampling can cause. We also have sequence a number of accessions which is, according to the known level of diversity of these species, supposed to be sufficient to be a good proxy to represent the species variability.

Secondly it depends on the used reference genomes which make a bias. In this study we have used the best reference genome that exist for the moment for rice. We are aware that, according to the phylogeny of rice, it would have been better to map our data against the African rice reference genome. We have chosen to use the Asian rice reference genome for several reasons. First because, the quality of this genome is much more better than the quality of the African rice reference genome in terms of annotations; and that is important in pan-genomic studies. Secondly because we though that, mapping the two species against a third one, give us a better view of their own differences. If we had chosen to map them against the African rice genome, then we might have expected to have better mapping results in the chosen species, and then, have a bigger bias for the second species compare to the first one. With the Asian rice as a reference genome, it is more or less like choosing an out group for a phylogeny study. However, we are conscious that, and for the future we are very interesting to remapped those data against the African rice genome. We hope that doing that, we could discover African rice specific genes or sequences.

Thirdly it depends on the strict definition on how to build the compartments (99% or 90% for the dispensable-genome for example). In this study we have chosen to stay on the initial definition of the pan-genome, meaning that being part of the dispensable-genome is possible with 99% of accessions having it. However, because sequencing technology is not 100% sure, it might be a better proxy to assign gene to the dispensable-genome if it is share by 90% of accessions, just like it has be done in other studies (REF !).

And finally it depends on the methodology you used to define if the gene is present or absent (based on depth of coverage, based on BLAST comparision, etc.). Pan-genomics analysis are mainly influenced by 6 aspects [?]:

1. The alignment algorithm and the parameters used to define similarity;
2. Phylogenetic resolution for the studied population;
3. Sampled individuals selected to represent the studied population;
4. Which model is used to estimate the number of new genes according to the number of genomes;
5. Type and quality of the annotation;
6. The level of comparison between the studied individuals (based for example on the sequence similarity or on presence/ absence profile of each gene independent the sequence similarity).

Pan-genomic analysis allows to determine the genomic diversity of a dataset, but also to predict how many extra genomes have to be add to characterize the whole pan-genome. For instance, extrapolation would be robust only if a high number of genomes is taken into account [?].

Vernikos et al [?] have inventoried 8 different methods dedicated to pan-genomic analyses:

- ORFsim: ORF alignment similarity;
- OG: method based on cluster of orthologs;
- Comb: combinatorial approach with successive addition of genomes;
- Gene freq: frequency of gene presence/ absence;
- Ref: generation of a reference pan-genome using a subset of individual;
- FSM: finite supragenome model;
- BMM: binomial mixture model;
- IMGM: infinitely many genes model.

Most of these methodologies were developed for bacterial or microorganism species, which are much more smaller and less complex than plants genomes (no polyploid, not so much of transposable elements for example). Those methodologies are also mostly based on cluster of orthologs, and get them on plant genome is a long process and we wanted something quicker. The tools developed for pan-genomic analysis are also mostly individual-centered. It means, we look to the difference of coverage between many loci inside one individual, and do it again on another one. It is good working, but, we thing that we should consider population. The final purpose of pan-genomic analysis is to do comparative analysis, so making it taking in account the population, was for us, a better way of doing a pan-genomic study. That is the reason why we developed a new method based neither on individual but on loci. With this method, we compare the difference of covering between individual for the same locus. Because the mapping coverage is different depending on the accessions, we beforehand have normalized the data. We can then look for outlier and so defined individual for which the gene is then considered to be absent.

We did not see any differences between the profile of the 12 chromosomes neither on the profile focus on exons nor on the profile focused on the remaining. This means there is no influence of the type of sequences (exon, CDS, UTR…) to the pan-genome compartment. This was confirm by the distribution of compartments along the chromosomes showing no significant profile, while Lu et al [Lu et al., 2015] found more PAV into the telomeric regions and few of them around the centromeres. However, we have noticed some hot-spot in the genome rich for one or the other compartment (faire un lien sur figure chr5, et ajouter une légende spécifique pour ça !!!). For further analysis we are interested in those regions to identify if one particular type of genes or sequences is behind these hot-spot. For example are those regions TE-rich? Have those regions a high GC percentage? We did not see any differences inside the *O. glaberrima* population, but in the wild one, we have detected two kinds of profiles. Indeed the core/pan ratio suggest that the whole accessions of *O. glaberrima* are extremely close to each other for the structure of their pan-genome. On the other hand, the *O. barthii* core/pan ratio suggest the accessions have more differences between them compare to intra *O. glaberrima*. It would be interesting to see if the two profiles we detect here, are based on the proximity of pan-genome structure of the accessions. If it is the case, that might be a new approach to getting a structural idea of one population. (???) In the present case, as already described in the results part, *O. barthii* species can be divided in two populations. We have tried to find a link between this two populations and the two profile without getting a clear answer. We concluded that it is neither a population effect nor the number of individuals tested which is important but rather the size of the initial population (meaning 121 for *O. glaberrima* instead of 23 for the population 1 of *O. barthii*). But still, it means that a minimal number of 50 individuals is better to have a robust profile in the case of African rices. That confirm that a pan-genomic study needs a minimal number of different accessions to get enough power to detect true biologically meaning information.

As the pan-genome of the cultivated species is smaller than the core-genome of the wild one, it confirms that there is a loss of diversity during the domestication process in term of presence/ absence variation. Which was not a surprise because it is what we already know from SNPs analysis. We also saw that the dispensable-genome of the cultivated is unexpectedly bigger than the wild one. Which means that in a certain way there is a new diversification inside the cultivated species after or during the domestication process which is stronger than the diversification forces acting on the wild species. The influence of human care one the cultivated genome could be the cause of the higher diversity we detect. By providing nutrients, water and, care against diseases and stress, humans might have influence the selection pressure on some part of the genome leading to a diversification of it. To confirm that we need first to investigate the genes/ sequences which are common to the dispensable-genome of the two species and, compare them to see their level of identity to determine a basal level for what is the differences between the two species. After that, we can compare what is dispensable-genome specific of each species and, try to get if, for example, the part of *O. glaberrima* is looking like pseudogenisation of what we detect in *O. barthii*. That would be a structural confirmation of the impact of human being on the genome. It would be also informative to have the pan-transcriptome of theses two species, to be able to say if the dispensable-part is functional or not. And if the level of expression are different when we look to the cultivated or the wild species. And then considering the effect of human on the functionality of genomes.

The fact that there are somes genes which move from core and dispensable-genome to the other between *O. barthii* and *O. glaberrima* means that in general when we go from the wild to the cultivated species we can observe a decrease of the number of alleles and core genes. But unexpectidly, we observe also an increase of the number of dispensable genes. The domestication process have promoted a few genes from the dispensable of the wild into the core-genome of the cultivated, but have also disavantaged a certain number of wild core genes to become dispensable into the cultivated species.

The fact that some genes move from the dispensable-genome to the core-genome between *O. barthii* and *O. glaberrima* can be easily explain by the process of selection. Indeed, by favoring some accessions, the core-genome of the selected pool can become bigger than the initial one. It might stay the same size if the choice of the accessions is done all around the diversity but, if it is bias (for agronomical reasons, for example) and we selected individuals who are closest to each other then, the core-genome we observe seems bigger and the real one (Figure taille core selection). On the opposite, the fact that genes move from the core-genome of the wild species to the dispensable-genome of the wild one is more surprising. This can not be explained by the selection but, probably it is link to domestication process on a certain number of generations. For the exact same reasons we have discussed above about the size of the pan-genome of the cultivated species which is smaller than the wild species. Through the domestication process, human care seems to have had an effect on the selection pressure for certain genes, and in extreme cases lead to the lose of these genes in certain genomes then becoming part of the dispensable-genome instead of core-genome. In the case of African rice, we know that only thousand generations have been necessary to create the cultivated species so, it would be interesting to do a modelisation trying to explain how genes of the core-genome can become dispensable-genome in a so short period of time. It would also be interessting to make some focus on genes that are know to be related to domestication to see if the selection pressure have changed for them. A good example might be the shattering gene (qSH1). This gene is know to have a SNP in the 5’ regulatory region leading to the loss of shattering in the cultivated species [?]. It would be interesting to see if the selection pressure for the gene is different for *O. barthii* and *O. glaberrima*. The proportion of dispensable-genome in the two species was as the opposite of what we expected it. We toughed the dispensable-genome, which is usually representing the variable part of the species would be bigger in the wild species because cultivated species are know to be very diversity poor. But, in the case of African rice at least, it is the exact opposite. We made this hypothesis because of what SNPs analysis has accustomed us to, but it seems it is different when you look in the genome structure. It might means that, the poor diversity we are able to detect, to see, is mostly due to SNPs but, cultivated species are not that poor of variability. In the case of *O. glaberrima* it is even richer than their wild relative. Perhaps this variability is not very usefull for agronomy, because these genes seems to be not relevant to do improvement of cultivation. But from a fundamental point of view this variability is there and by a deeper analysis of it, might become usefull. The same idea is indicated by the ratio corep̃an-genome. As presented in bacteria by Caputo et al [Caputo et al., 2015] a ratio core/ pan-genome higher than 90% is indicating a high rate of conservation between the individual of the sample. In our case, the core/ pan-genome ratio is equal to 98.15% for *O. barthii* meaning that the individuals are all really close to each other. Surprisingly for *O. glaberrima*, it decreases to 86.44% meaning that the cultivated individuals are more divergent to each other than the wild together.

With the GO, we showed that it is not necessary for the all members of one gene family to belong to the same compartment, which might contribute to the hypothesis that members of the same gene family can complement each other. We also confirm the movement from one compartment to another between the two species. We retrieved the gene coming from the core to the dispensable during the domestication. By this way we also find out genes which are missing in the pan-genome of the cultivated species. Among them we have found the GO in response to stress or stimulus. These one could be easily explain why they are absent of *O. glaberrima*, because of the input made by human in cultivation. As agriculture brings defenses against pathogenes and challenging conditions, the organism does not need to be able to protect himself against it. This is in correlation with the hypothesis at, with domestication process and human being activities, plants are assisted and do not need certain genes to survive anymore. This theory was developped by Loss et al [Loss, 2012] describing the wild species with a lot of genes to be able to anwser for a lot of stress, and the cultivated one, who loss genes because human being provides defenses or nutrients to compensate this gene loss.

In this study we performed the first pan-genome analyses on African rices. It is also the first study comparing both cultivated species and its wild relative by a pan-genome approach. Doing it we confirmed the loss of diversity during the domestication process because the total number of genes of the cultivated species is lower than the core-genome of the wild one. Despite it the cultivated species developped a new diversity in term of dispensable genes. This diversity seems to be really active as it is higher than the dispensable diversity of the wild species. Our hypothesis is that domestication process act on diversity at two step. First the initial selection from the *O. barthii* reduces the diversity as we choose only prefered individuals. Then, as the domestication process long for some generations - in our case about 1,000- the selection pressure provokes either a drop of it or a negative agronomic one act to finally create new diversity inside the new species to give the actual cultivated species.

## Acknowledgments

Authors wants to thanks Roland Akakpo for his R script to analyse the GO.

---

**Table S1.** Sequencing coverage on the *Oryza glaberrima* and *Oryza barthii* accessions

|  | *Oryza glaberrima* | *Oryza barthii* |
| --- | --- | --- |
| mini coverage | 21x | 23x |
| maxi coverage | 52x | 48x |
| mean coverage | 37x | 33x |

**Table S2.**
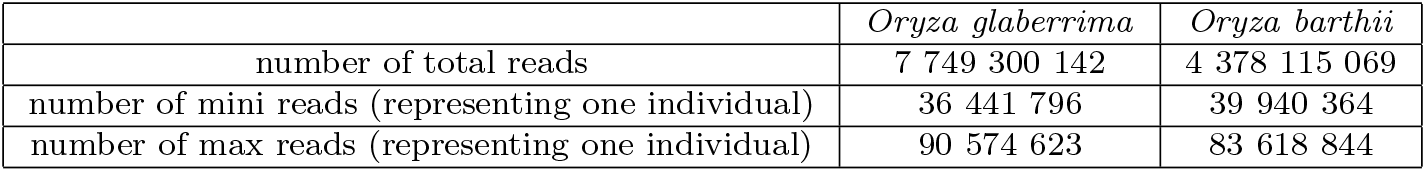
Number of reads on the *Oryza glaberrima* and *Oryza barthii* accessions

**Figure S1.**
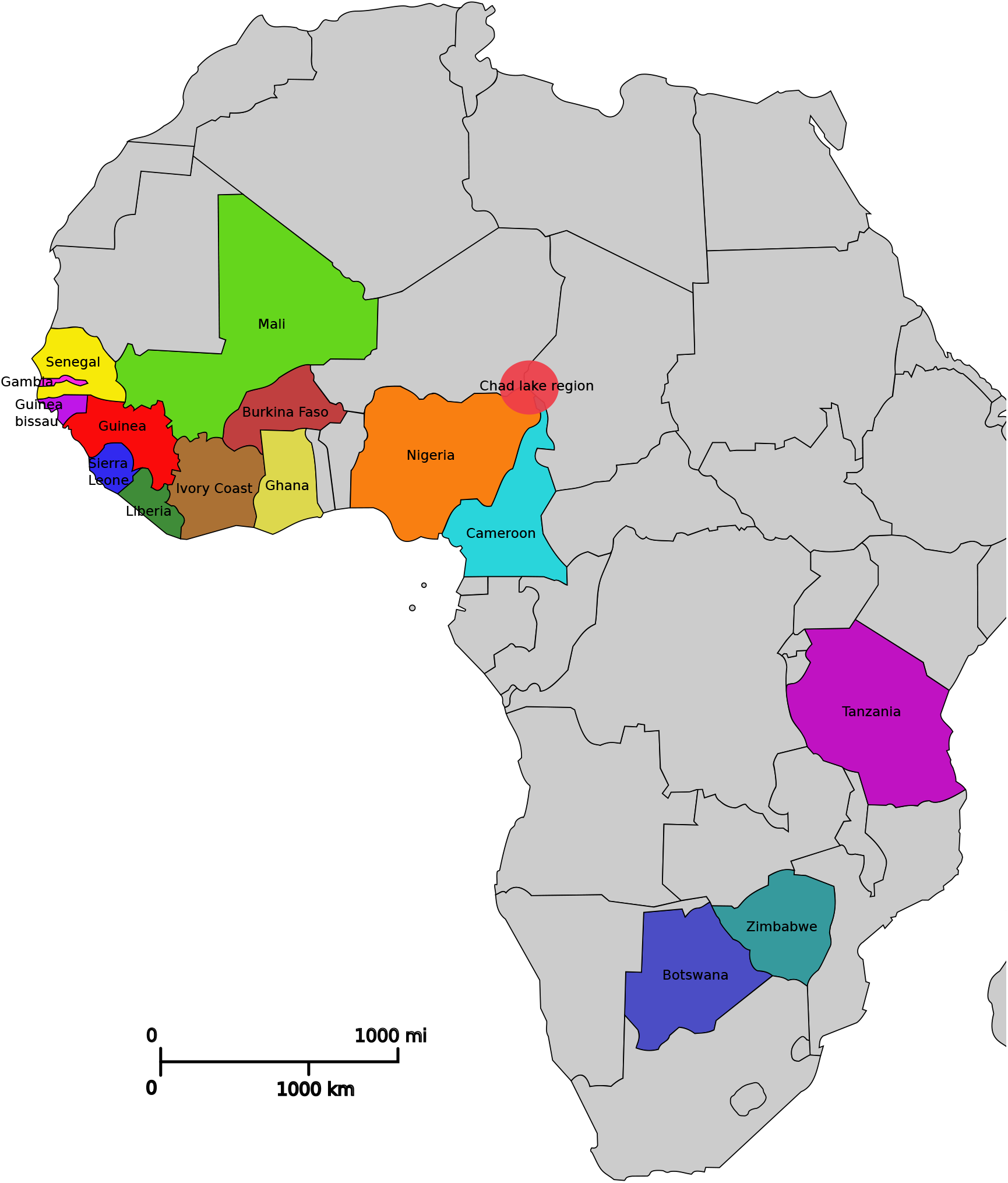
Geographical distribution of the accessions used in this analysis

**Figure S2.**
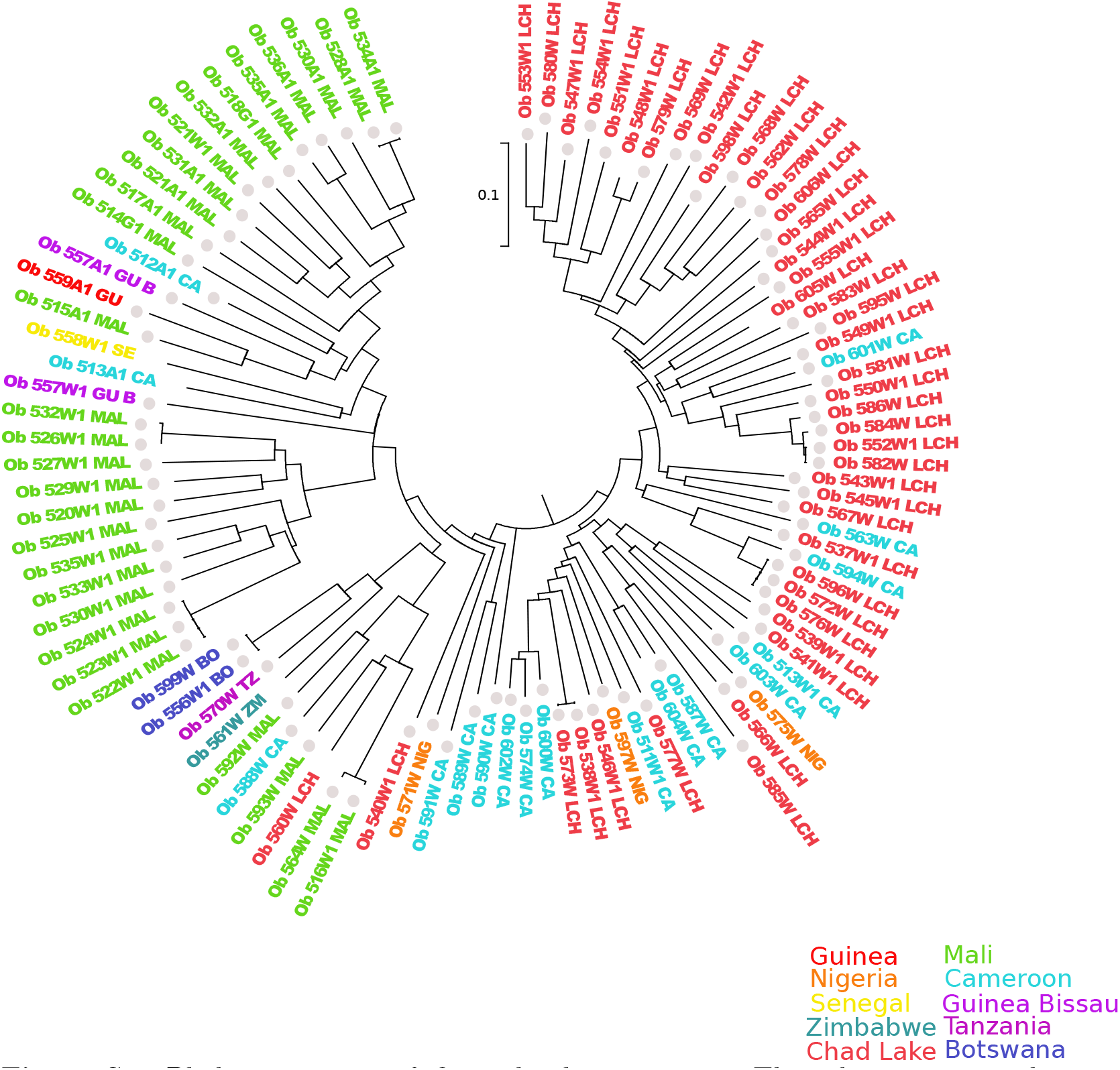
Phylogenetic tree of *Oryza barthii* accessions. The colors represent the countries just as in Fig. S1. The circles represent the accessions in this analysis. Adapted from Orjuela et al [Orjuela et al., 2014].

**Figure S3.**
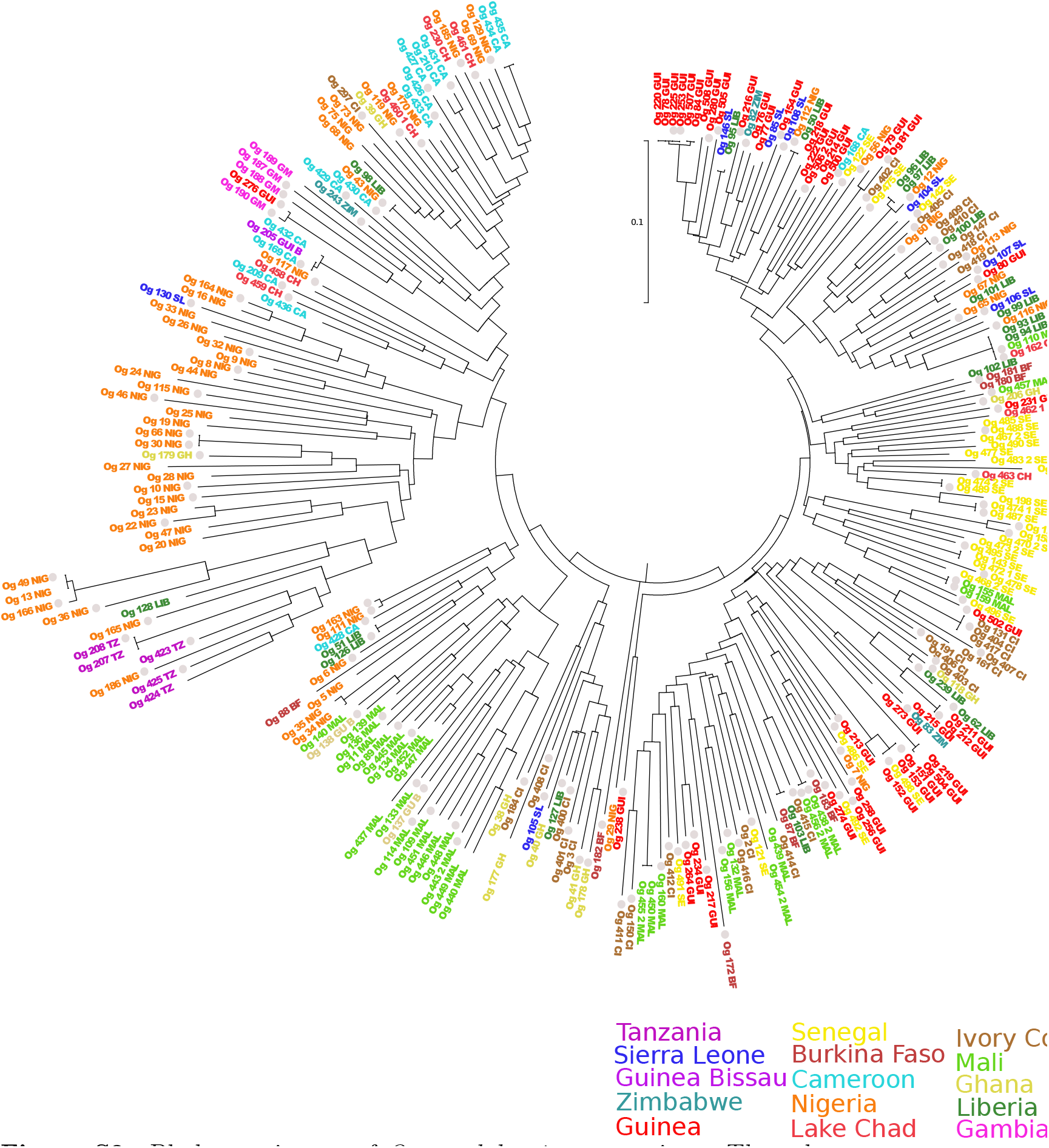
Phylogenetic tree of *Oryza glaberrima* accessions. The colors represent the countries just as in Fig. S1. The circles represent the accessions in this analysis. Adapted from Orjuela et al [Orjuela et al., 2014].

**Figure S4.**
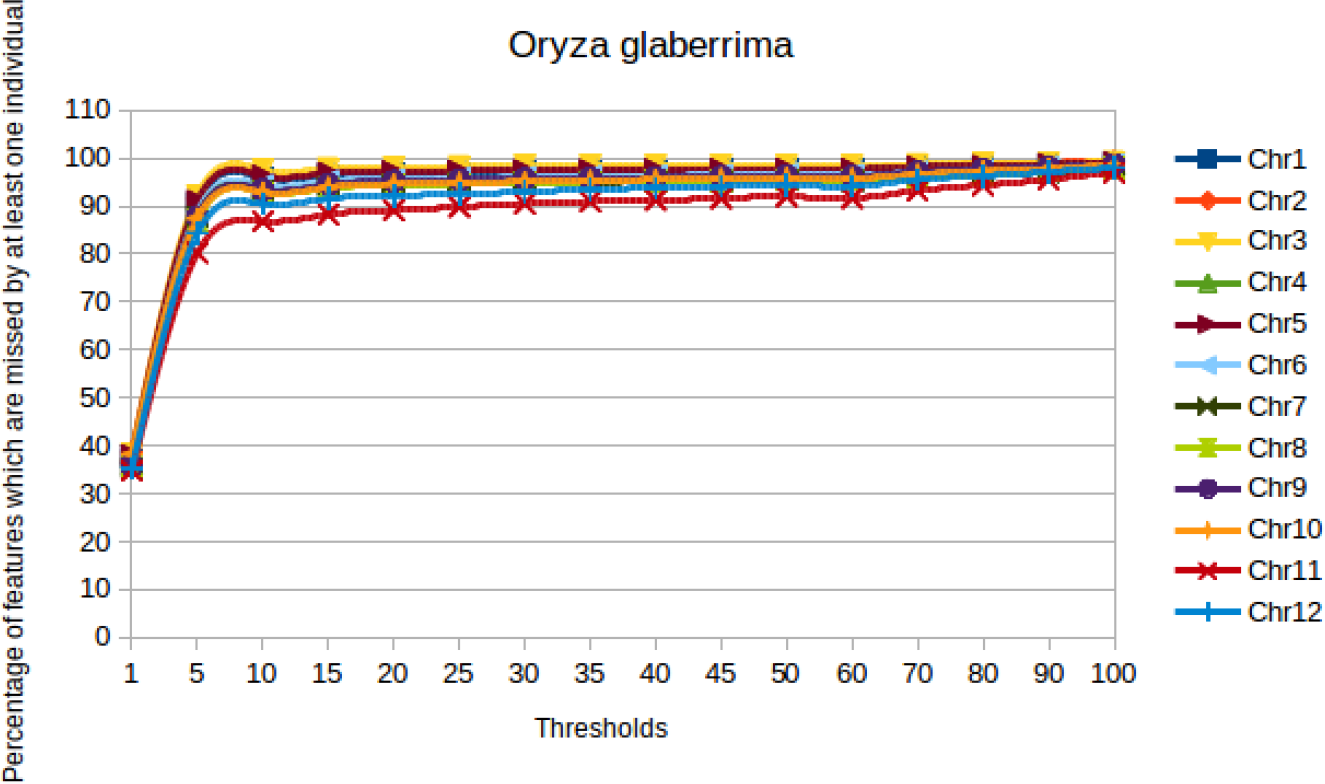
Percentage of annotation features with at least one outlier among the *O. glaberrima* species

**Figure S5.**
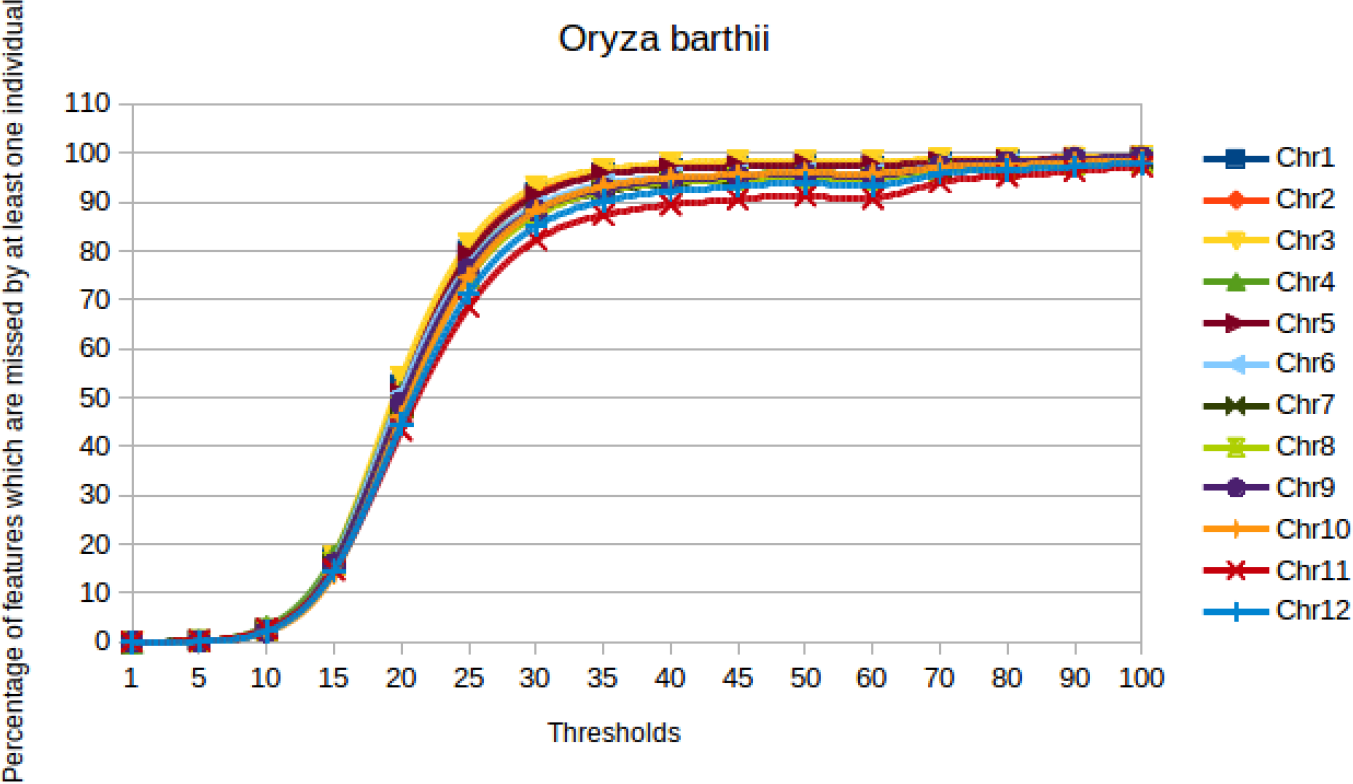
Percentage of annotation features with at least one outlier among the *O. barthii* species

**Figure S6.**
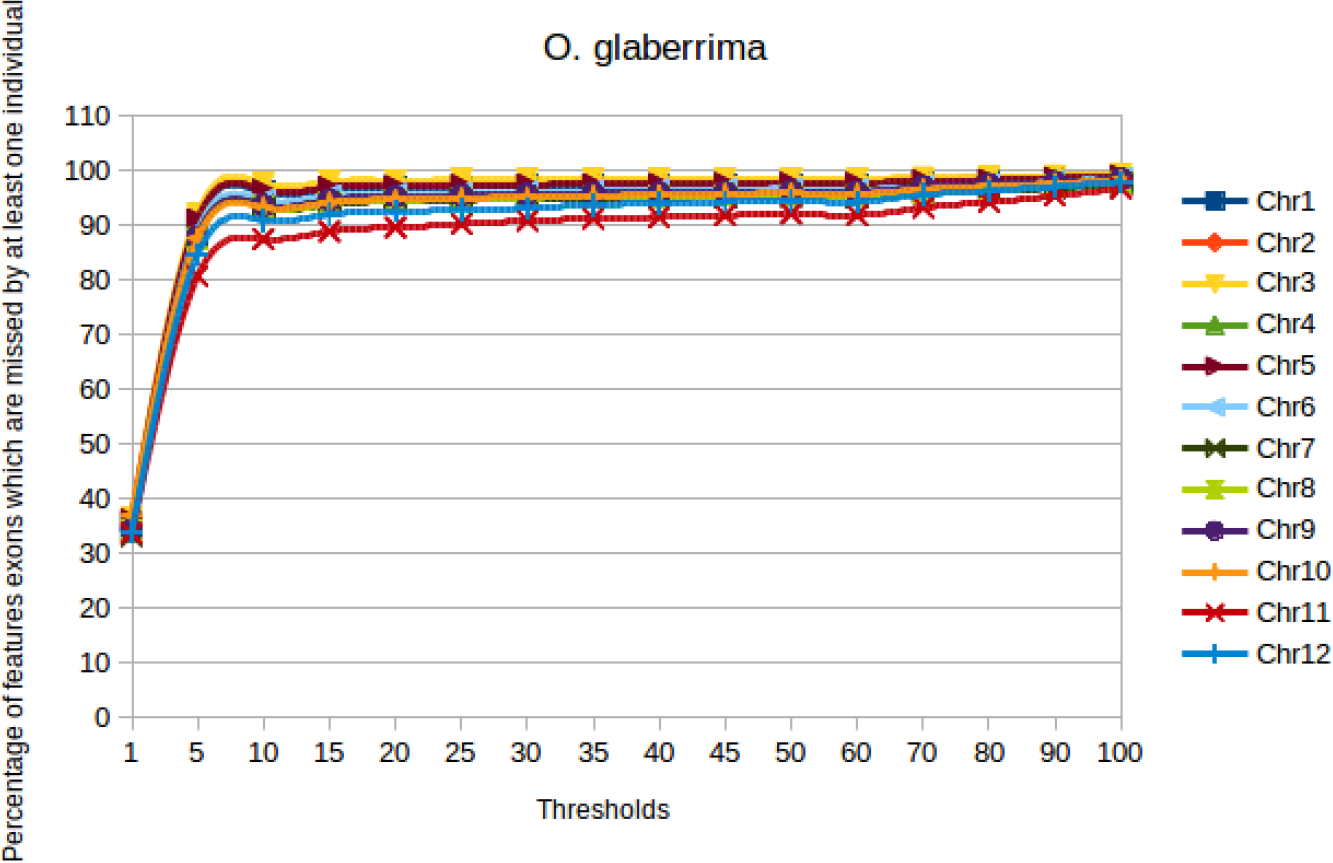
Percentage of exons features with at least one outlier among the *O. glaberrima* species

**Figure S7.**
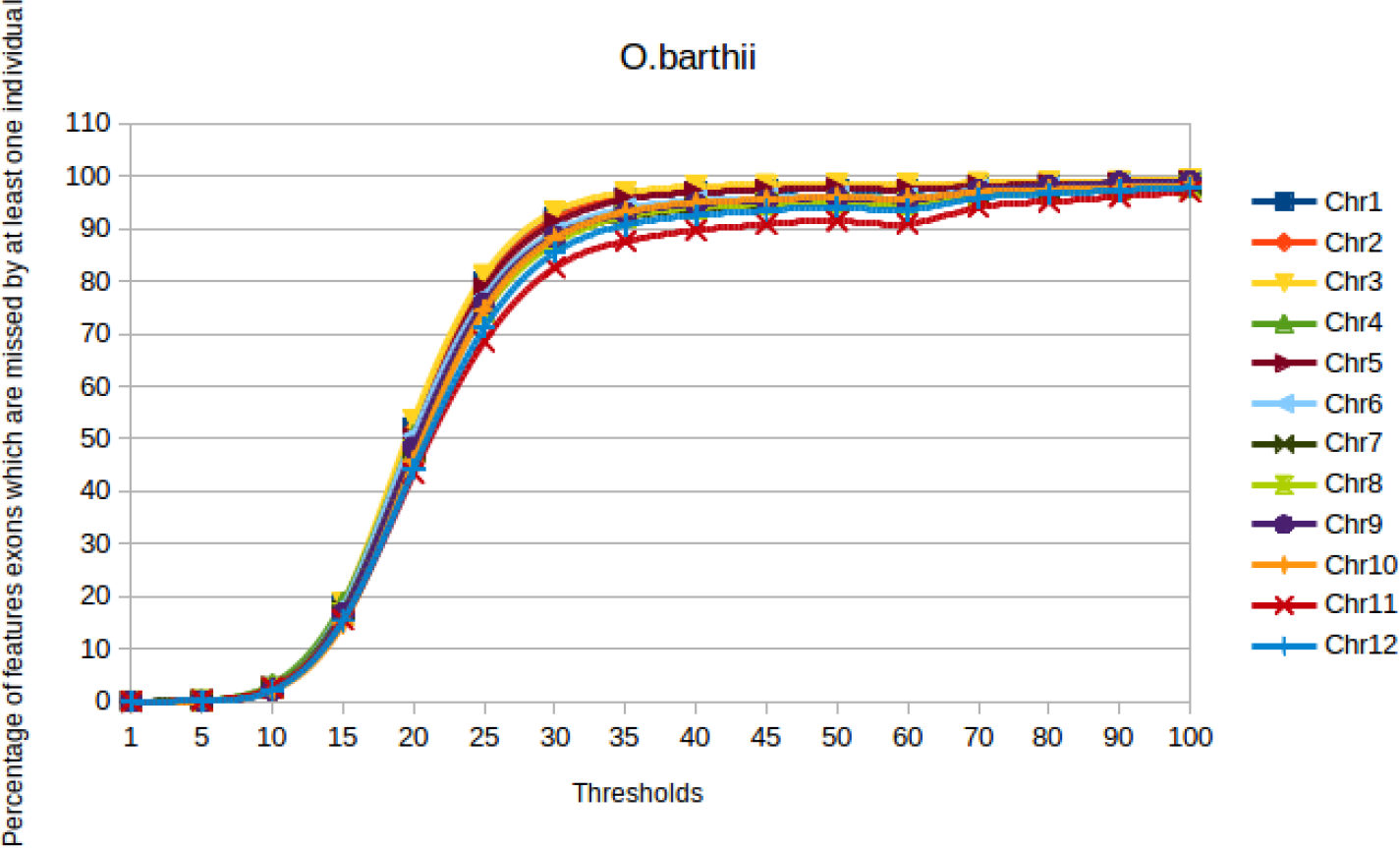
Percentage of exons features with at least one outlier among the *O. barthii* species

**Figure S8.**
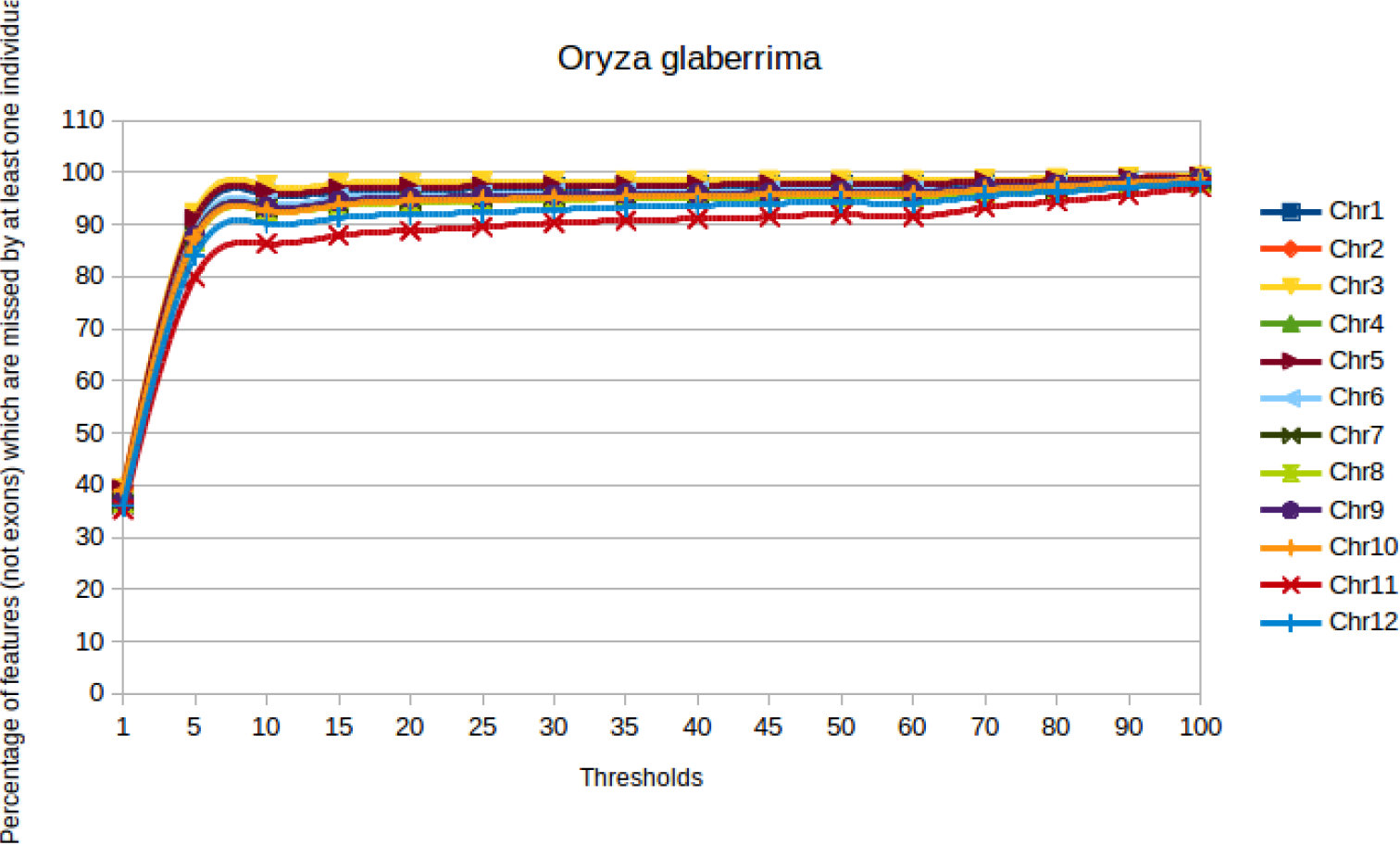
Percentage of not exons features with at least one outlier among the *O. glaberrima* species

**Figure S9.**
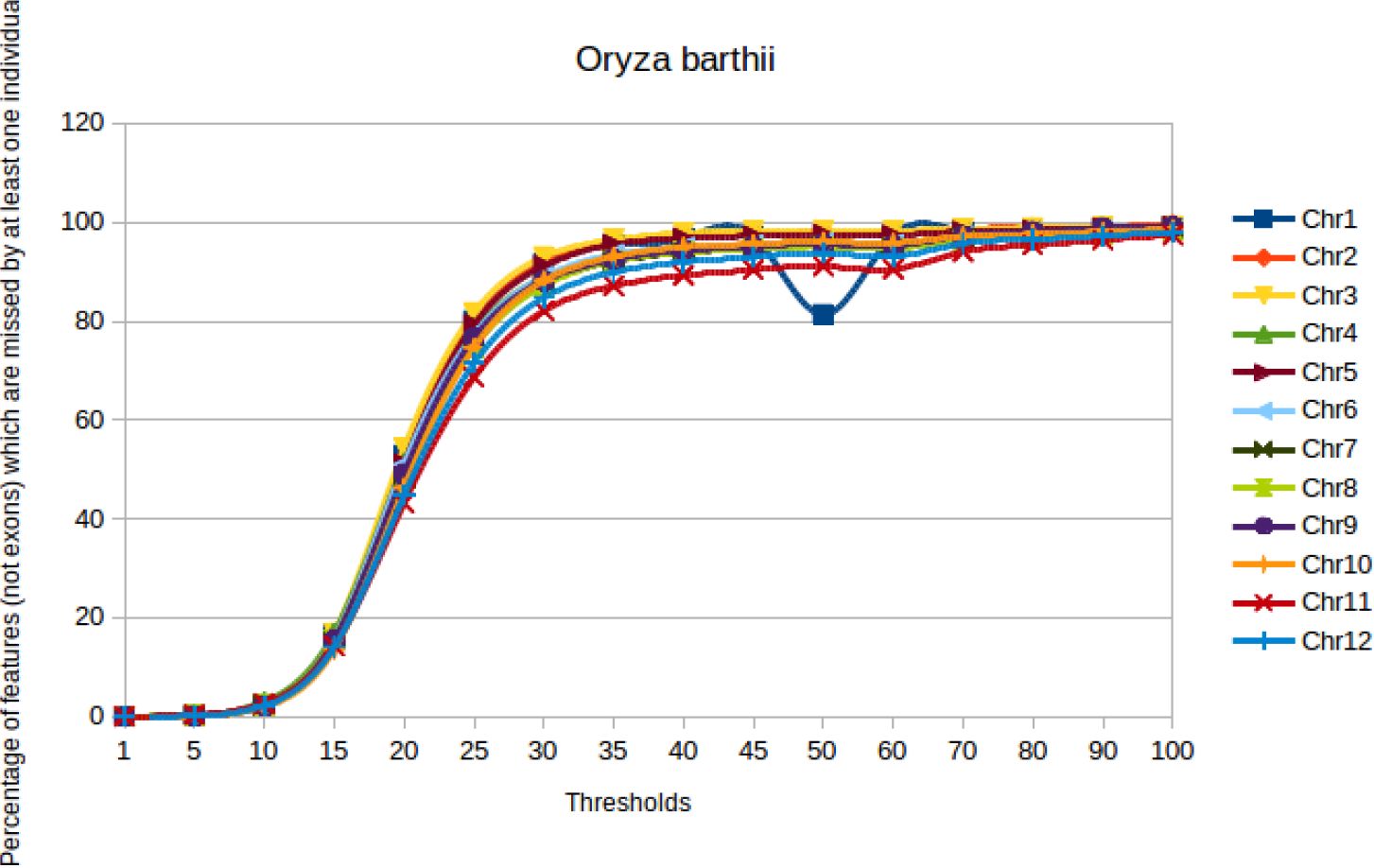
Percentage of not exons features with at least one outlier among the *O. barthii* species

**Figure S10.**
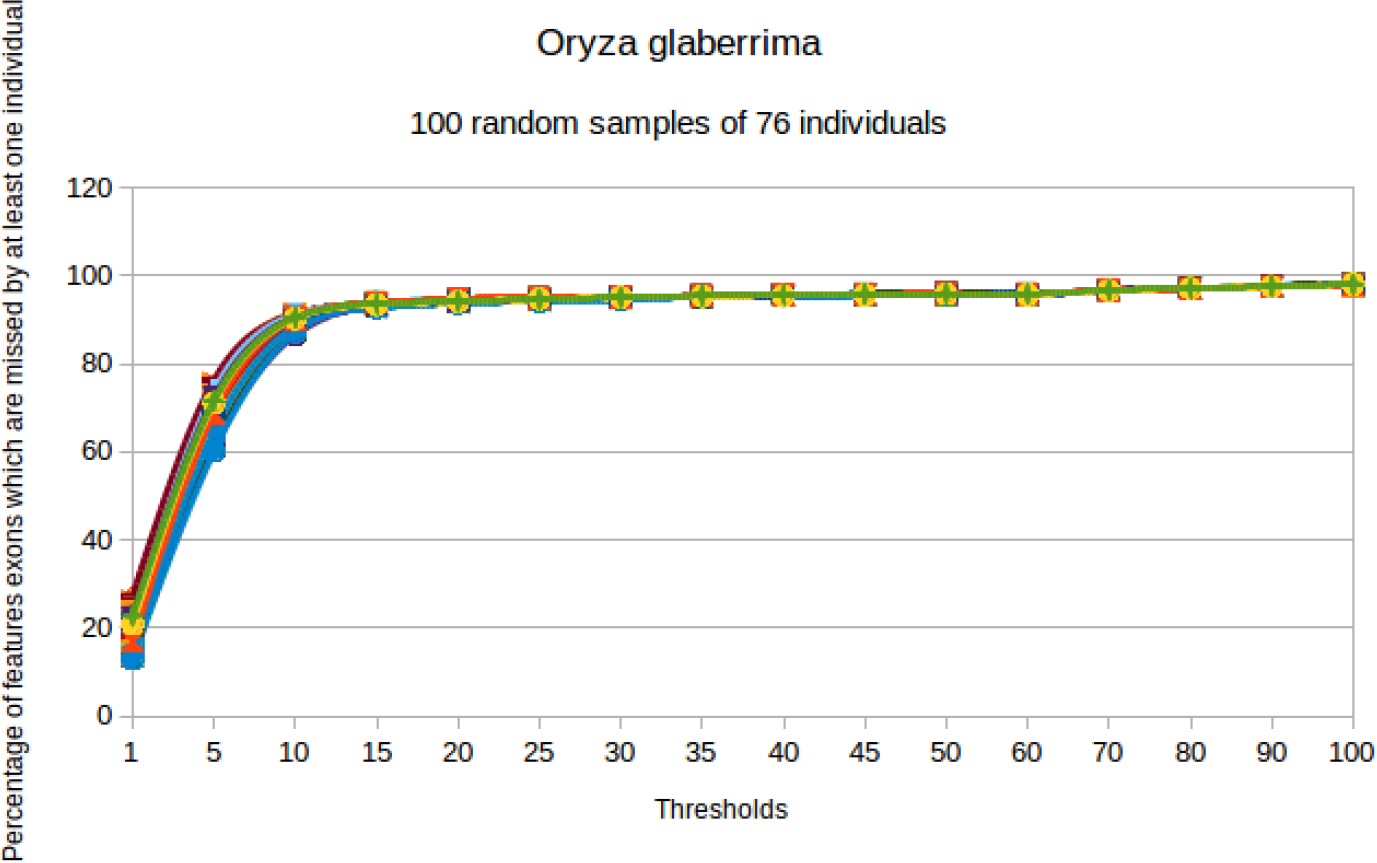
Hundred bootstrap of 76 individuals from 120 accessions of *O. glaberrima*

**Figure S11.**
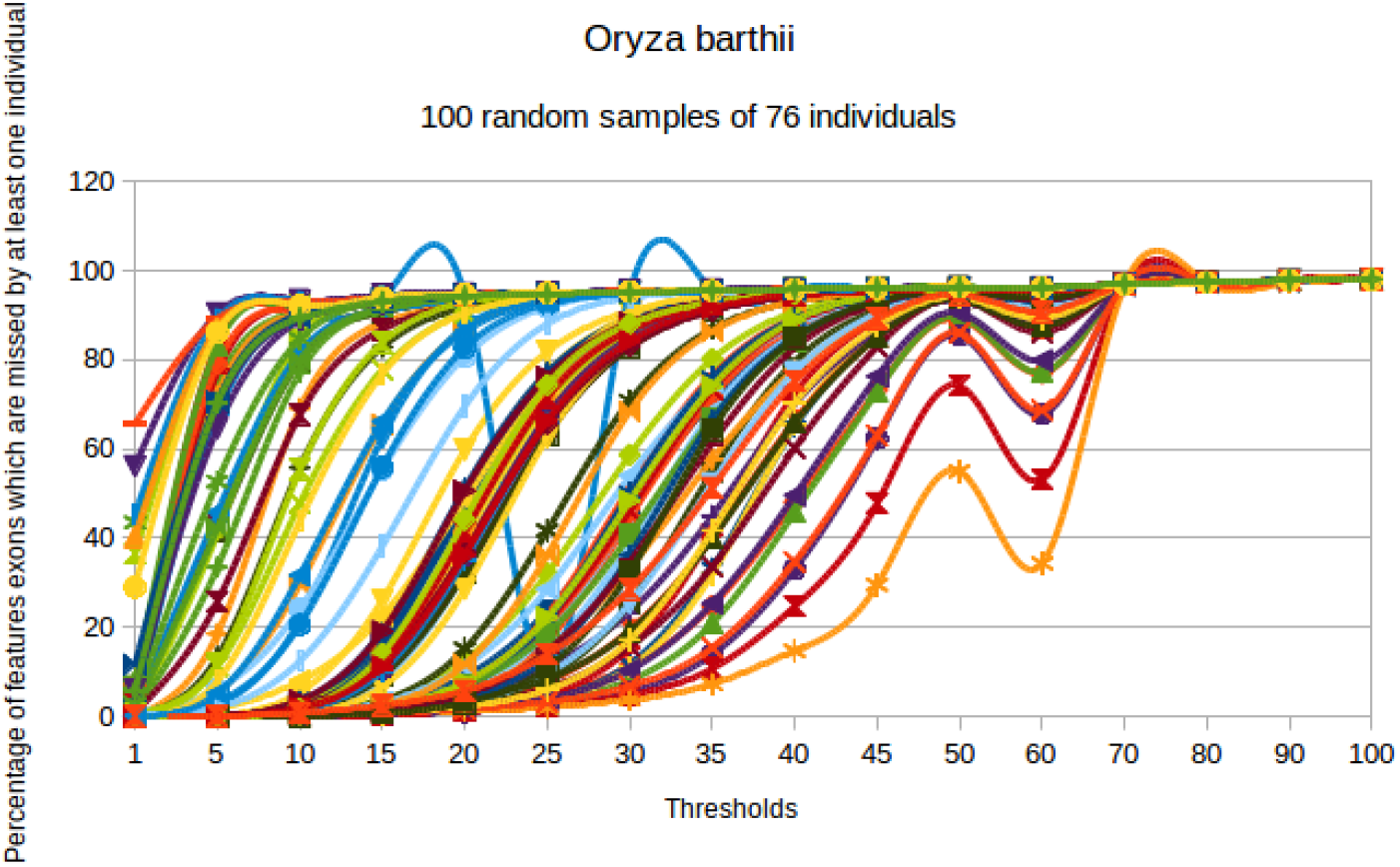
Hundred bootstrap of 76 individuals from 76 accessions of *O. barthii*

**Figure S12.**
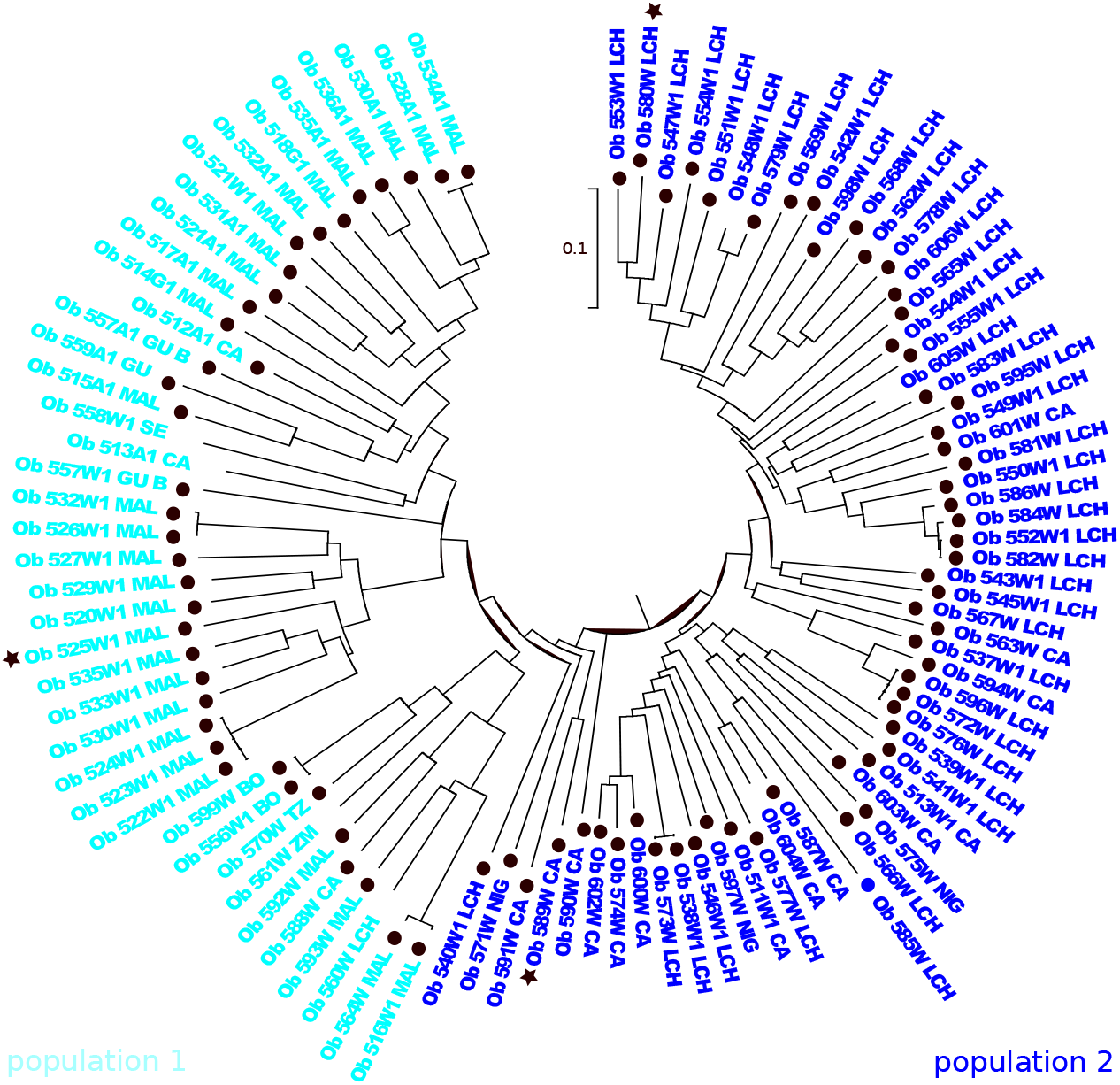
Phylogenetic tree of *O. barthii* with the population 1 accessions on cyan and the population 2 accessions on navy blue. Adapted from Orjuela et al [Orjuela et al., 2014]

**Figure S13.**
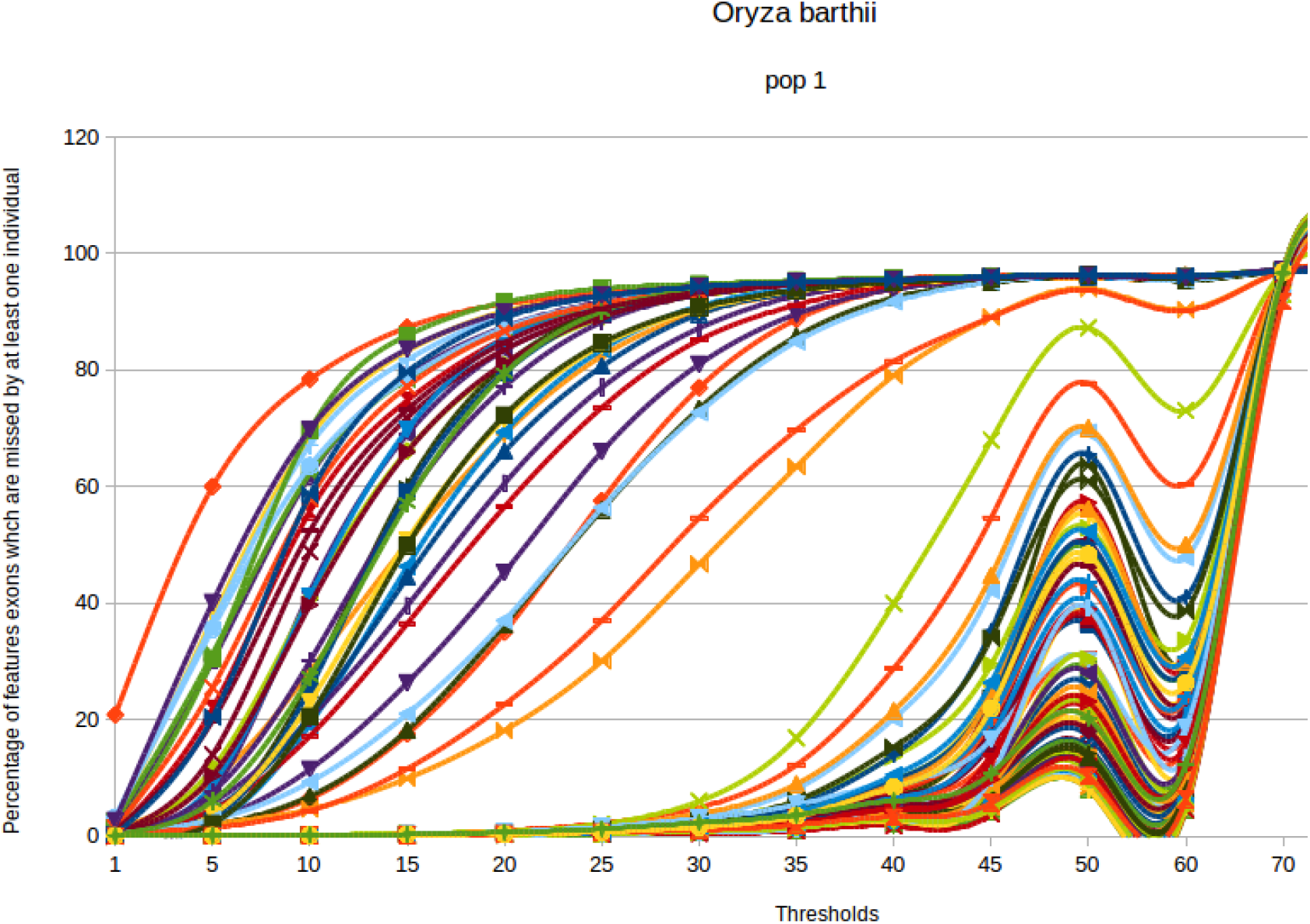
Hundred bootstrap of 23 individuals from 23 accessions of population 1 of *O. barthii*

**Figure S14.**
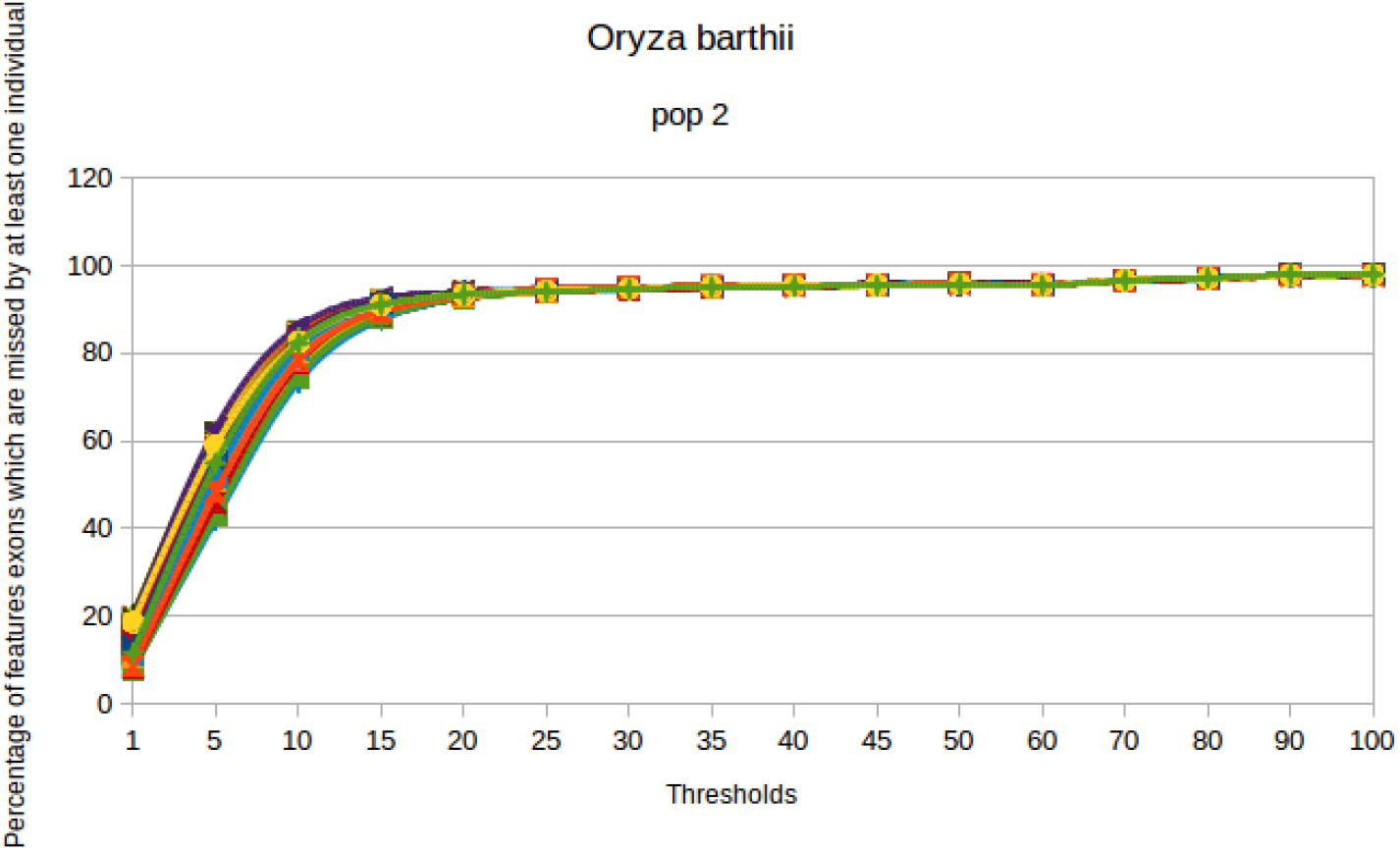
Hundred bootstrap of 51 individuals from 51 accessions of population 2 of *O. barthii*

**Figure S15.**
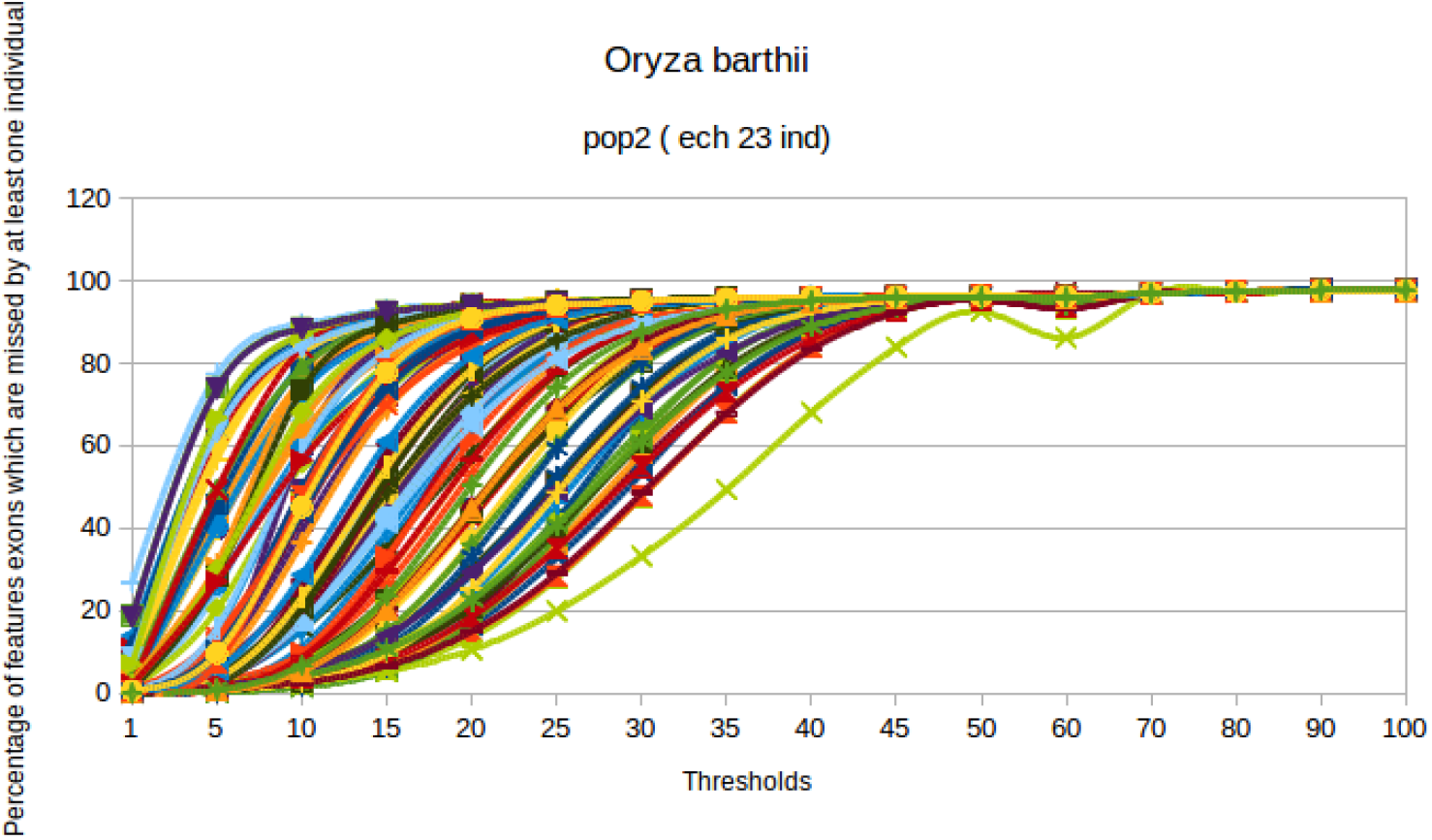
Hundred bootstrap of 23 individuals from 51 accessions of population 2 of *O. barthii*

**Figure S16.**
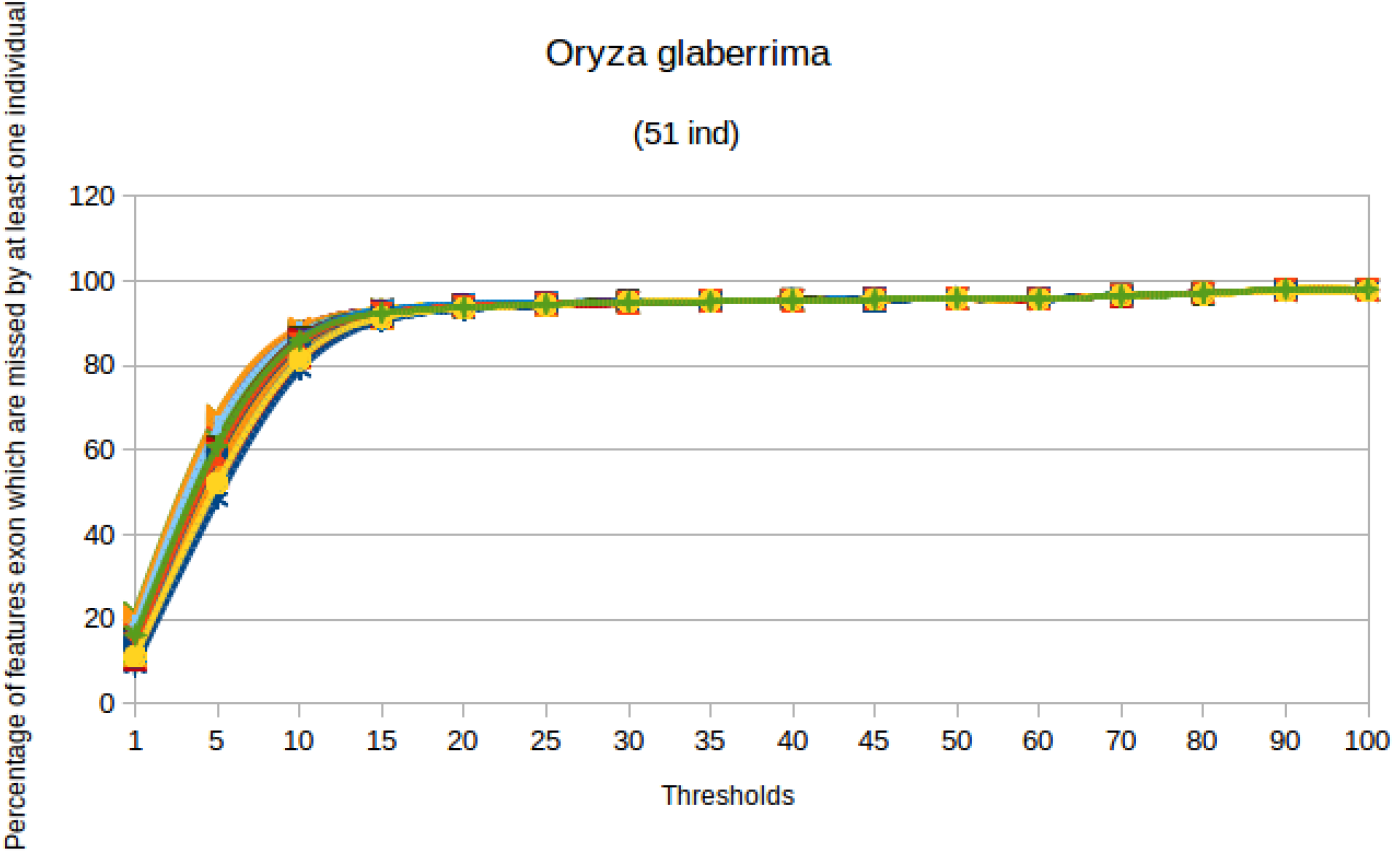
Hundred bootstrap of 51 individuals from 120 accessions of *O. glaberrima*

**Figure S17.**
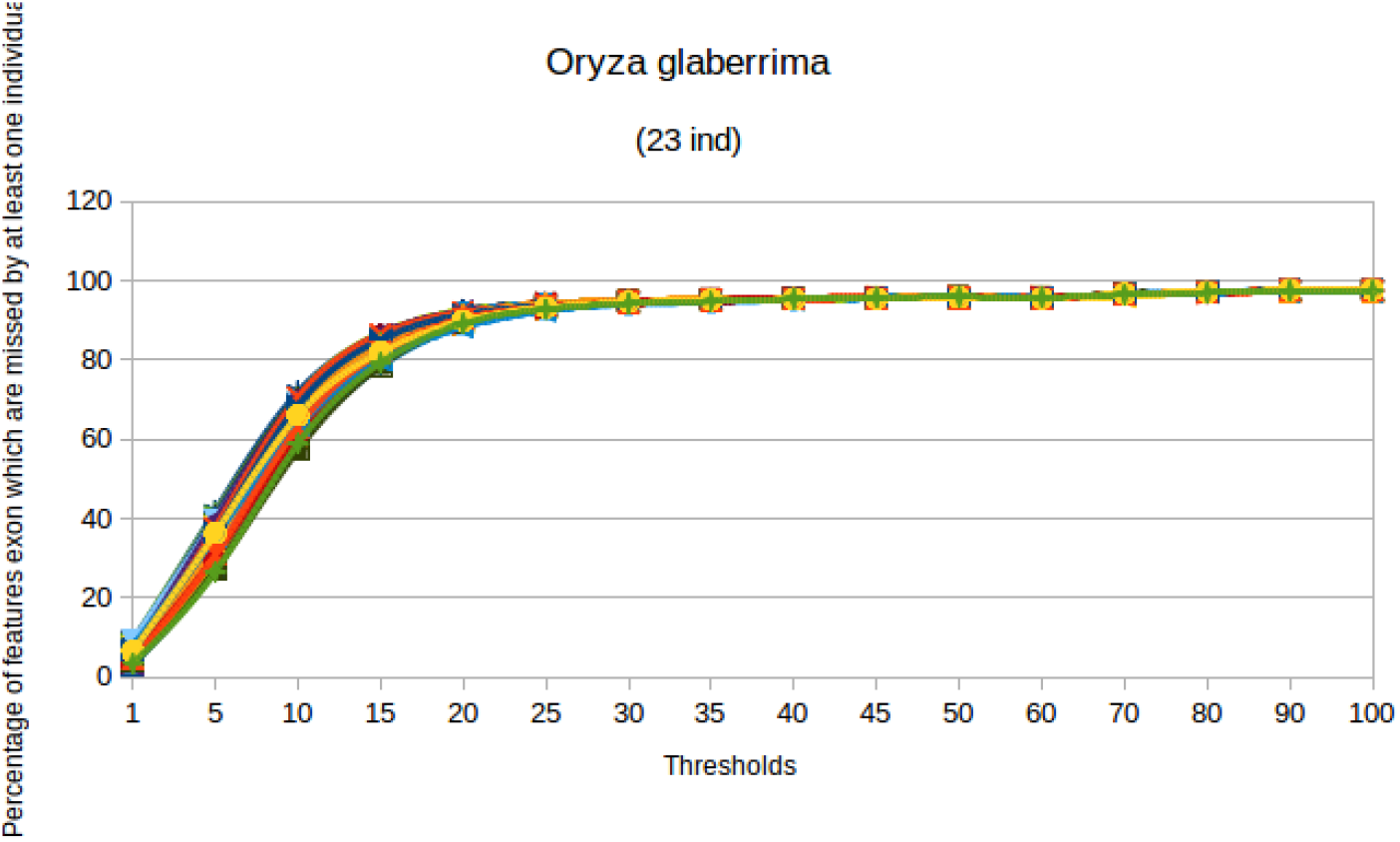
Hundred bootstrap of 23 individuals from 120 accessions of *O. glaberrima*

## References

Cub, 2017. (2017). Genome footprints of the rise and fall of African rice cultivation. under Review, 1:1–10.

Aherfi et al., 2014. Aherfi, S., La Scola, B., Pagnier, I., Raoult, D., and Colson, P. (2014). The expanding family Marseilleviridae. Virology, 467:27–37.

Alexa and Rahnenfuhrer, 2016. Alexa, A. and Rahnenfuhrer, J. (2016). topGO: Enrichment Analysis for Gene Ontology. R package version 2.24.0.

Boussaha et al., 2015. Boussaha, M., Esquerré, D., Barbieri, J., Djari, A., Pinton, A., Letaief, R., Salin, G., Escudié, F., Roulet, A., Fritz, S., Samson, F., Grohs, C., Bernard, M., Klopp, C., Boichard, D., and Rocha, D. (2015). Genome-Wide Study of Structural Variants in Bovine Holstein, Montbéliarde and Normande Dairy Breeds. Plos One, 10(8):e0135931.

Caputo et al., 2015. Caputo, A., Merhej, V., Georgiades, K., Fournier, P.-E., Croce, O., Robert, C., and Raoult, D. (2015). Pan-genomic analysis to redefine species and subspecies based on quantum discontinuous variation: the Klebsiella paradigm. Biology Direct, 10(1):55.

Cheung et al., 2009. Cheung, F., Trick, M., Drou, N., Lim, Y. P., Park, J.-Y., Kwon, S.-J., Kim, J.-A., Scott, R., Pires, J. C., Paterson, A. H., Town, C., and Bancroft, I. (2009). Comparative Analysis between Homoeologous Genome Segments of Brassica napus and Its Progenitor Species Reveals Extensive Sequence-Level Divergence. the Plant Cell Online, 21(7):1912–1928.

Chia et al., 2012. Chia, J.-M., Song, C., Bradbury, P. J., Costich, D., de Leon, N., Doebley, J., Elshire, R. J., Gaut, B., Geller, L., Glaubitz, J. C., Gore, M., Guill,K. E., Holland, J., Hufford, M. B., Lai, J., Li, M., Liu, X., Lu, Y., McCombie, R., Nelson, R., Poland, J., Prasanna, B. M., Pyhäjärvi, T., Rong, T., Sekhon, R. S., Sun, Q., Tenaillon, M. I., Tian, F., Wang, J., Xu, X., Zhang, Z., Kaeppler, S. M., Ross-Ibarra, J., McMullen, M. D., Buckler, E. S., Zhang, G., Xu, Y., and Ware, D. (2012). Maize HapMap2 identifies extant variation from a genome in flux. Nature genetics, 44(7):803–7.

Conte et al., 2008a. Conte, M. G., Gaillard, S., Droc, G., and Perin, C. (2008a). Phylogenomics of plant genomes: a methodology for genome-wide searches for orthologs in plants. BMC genomics, 9:183.

Conte et al., 2008b. Conte, M. G., Gaillard, S., Lanau, N., Rouard, M., and Périn, C. (2008b). GreenPhylDB: A database for plant comparative genomics. Nucleic Acids Research, 36(SUPPL. 1):991–998.

Gan et al., 2011. Gan, X., Stegle, O., Behr, J., Steffen, J. G., Drewe, P., Hildebrand, K. L., Lyngsoe, R., Schultheiss, S. J., Osborne, E. J., Sreedharan, V. T., Kahles, A., Bohnert, R., Jean, G., Derwent, P., Kersey, P., Belfield, E. J., Harberd, N. P., Kemen, E., Toomajian, C., Kover, P. X., Clark, R. M., Rätsch, G., and Mott, R. (2011). Multiple reference genomes and transcriptomes for Arabidopsis thaliana. Nature, 477(7365):419–23.

Kawahara et al., 2013. Kawahara, Y., de la Bastide, M., Hamilton, J. P., Kanamori, H., McCombie, W. R., Ouyang, S., Schwartz, D. C., Tanaka, T., Wu, J., Zhou, S., Childs, K. L., Davidson, R. M., Lin, H., Quesada-Ocampo, L., Vaillancourt, B., Sakai, H., Lee, S. S., Kim, J., Numa, H., Itoh, T., Buell, C. R., and Matsumoto, T. (2013). Improvement of the Oryza sativa Nipponbare reference genome using next generation sequence and optical map data. Rice (New York, N.Y.), 6(1):4.

Laing et al., 2010. Laing, C., Buchanan, C., Taboada, E. N., Zhang, Y., Kropinski, A., Villegas, A., Thomas, J. E., and Gannon, V. P. J. (2010). Pan-genome sequence analysis using Panseq: an online tool for the rapid analysis of core and accessory genomic regions. BMC bioinformatics, 11:461.

Li and Durbin, 2009. Li, H. and Durbin, R. (2009). Fast and accurate short read alignment with Burrows-Wheeler transform. Bioinformatics (Oxford, England), 25(14):1754–60.

Li et al., 2009. Li, H., Handsaker, B., Wysoker, A., Fennell, T., Ruan, J., Homer, N., Marth, G., Abecasis, G., and Durbin, R. (2009). The Sequence Alignment/Map format and SAMtools. Bioinformatics, 25(16):2078–2079.

Li et al., 2010. Li, R., Li, Y., Zheng, H., Luo, R., Zhu, H., Li, Q., Qian, W., Ren, Y., Tian, G., Li, J., Zhou, G., Zhu, X., Wu, H., Qin, J., Jin, X., Li, D., Cao, H., Hu, X., Blanche, H., Cann, H., Zhang, X., Li, S., Bolund, L., Kristiansen, K., Yang, H., Wang, J., and Wang, J. (2010). Building the sequence map of the human pan-genome. Nature biotechnology, 28(1):57–63.

Loss, 2012. Loss, A. G. (2012). The Black Queen Hypothesis: Evolution of Dependencies through. 3(2):1–7.

Lu et al., 2015. Lu, F., Romay, M. C., Glaubitz, J. C., Bradbury, P. J., Elshire, R. J., Wang, T., Li, Y. Y., Li, Y. Y., Semagn, K., Zhang, X., Hernandez, A. G., Mikel,M. A., Soifer, I., Barad, O., and Buckler, E. S. (2015). High-resolution genetic mapping of maize pan-genome sequence anchors. Nature Communications, 6:6914.

Lukjancenko et al., 2012. Lukjancenko, O., Ussery, D. W., and Wassenaar, T. M. (2012). Comparative Genomics of Bifidobacterium, Lactobacillus and Related Probiotic Genera. Microbial Ecology, 63(3):651–673.

Mckenna et al., 2010. Mckenna, A., Hanna, M., Banks, E., Sivachenko, A., Cibulskis, K., Kernytsky, A., Garimella, K., Altshuler, D., Gabriel, S., Daly, M., DePristo,M. a., and Others (2010). The Genome Analysis Toolkit: a MapReduce framework for analyzing next-generation DNA sequencing data. Genome research, 20(9):1297–1303.

Mcnally et al., 2009. Mcnally, K. L., Childs, K. L., Bohnert, R., Davidson, R. M., Zhao, K., Ulat, V. J., Zeller, G., Clark, R. M., Hoen, D. R., Bureau, T. E., Stokowski, R., Ballinger, D. G., Frazer, K. A., Cox, D. R., Padhukasahasram, B., Bustamante, C. D., Weigel, D., Mackill, D. J., Buell, C. R., Leung, H., Leach, J. E., Bruskiewich, R. M., and Ra, G. (2009). Genomewide SNP variation reveals relationships among landraces and modern varieties of rice. pages 1–6.

Monat et al., 2015. Monat, C., Tranchant-Dubreuil, C., Kougbeadjo, A., Farcy, C., Ortega-Abboud, E., Amanzougarene, S., Ravel, S., Agbessi, M., Orjuela-Bouniol, J., Summo, M., and Sabot, F. (2015). TOGGLE: toolbox for generic NGS analyses. BMC Bioinformatics, 16.

Montenegro et al., 2017. Montenegro, J. D., Golicz, A. A., Bayer, P. E., Hurgobin, B., Lee, H., Chan, C.-K. K., Visendi, P., Lai, K., Doležel, J., Batley, J., and Edwards, D. (2017). The pangenome of hexaploid bread wheat. The Plant Journal.

Nabholz et al., 2014. Nabholz, B., Sarah, G., Sabot, F., Ruiz, M., Adam, H., Nidelet, S., Ghesquière, A., Santoni, S., David, J., and Glémin, S. (2014). Transcriptome population genomics reveals severe bottleneck and domestication cost in the African rice (O. glaberrima). Molecular Ecology, 23(9):n/a–n/a.

Nieselt, 2015. Nieselt, K. (2015). Pan-Tetris: an interactive visualisation for. BioVis 2015(Suppl 11):1–11.

Orjuela et al., 2014. Orjuela, J., Sabot, F., Chéron, S., Vigouroux, Y., Adam, H., Chrestin, H., Sanni, K., Lorieux, M., and Ghesquière, A. (2014). An extensive analysis of the African rice genetic diversity through a global genotyping. TAG. Theoretical and applied genetics. Theoretische und angewandte Genetik, 127(10):2211–23.

Quinlan, 2014. Quinlan, A. R. (2014). BEDTools: The Swiss-Army Tool for Genome Feature Analysis., volume 47.

Rouard et al., 2011. Rouard, M., Guignon, V., Aluome, C., Laporte, M. A., Droc, G., Walde, C., Zmasek, C. M., Périn, C., and Conte, M. G. (2011). GreenPhylDB v2.0: Comparative and functional genomics in plants. Nucleic Acids Research, 39(SUPPL.1):1095–1102.

Sarovich and Price, 2014. Sarovich, D. S. and Price, E. P. (2014). SPANDx: a genomics pipeline for comparative analysis of large haploid whole genome re-sequencing datasets. BMC research notes, 7(1):618.

Snipen and Ussery, 2010. Snipen, L. and Ussery, D. W. (2010). Standard operating procedure for computing pangenome trees. Standards in genomic sciences, 2(1):135–141.

Tettelin et al., 2005. Tettelin, H., Masignani, V., Cieslewicz, M. J., Donati, C., Medini, D., Ward, N. L., Angiuoli, S. V., Crabtree, J., Jones, A. L., Durkin, a. S., Deboy, R. T., Davidsen, T. M., Mora, M., Scarselli, M., Margarit y Ros, I., Peterson, J. D., Hauser, C. R., Sundaram, J. P., Nelson, W. C., Madupu, R., Brinkac, L. M., Dodson, R. J., Rosovitz, M. J., Sullivan, S. a., Daugherty, S. C., Haft, D. H., Selengut, J., Gwinn, M. L., Zhou, L., Zafar, N., Khouri, H., Radune,D., Dimitrov, G., Watkins, K., O’Connor, K. J. B., Smith, S., Utterback, T. R., White, O., Rubens, C. E., Grandi, G., Madoff, L. C., Kasper, D. L., Telford, J. L., Wessels, M. R., Rappuoli, R., and Fraser, C. M. (2005). Genome analysis of multiple pathogenic isolates of Streptococcus agalactiae: implications for the microbial “pan-genome”. Proceedings of the National Academy of Sciences of the United States of America, 102(39):13950–5.

Tettelin et al., 2008. Tettelin, H., Riley, D., Cattuto, C., and Medini, D. (2008). Comparative genomics: the bacterial pan-genome. Current Opinion in Microbiology, 11(5):472–477.

Thakur and Guttman, 2016. Thakur, S. and Guttman, D. S. (2016). A de-novo genome analysis pipeline (DeNoGAP) For Large-Scale Comparative Prokaryotic Genomics Studies. BMC bioinformatics, pages 1–18.

Upadhyaya et al., 2015. Upadhyaya, N. M., Garnica, D. P., Karaoglu, H., Sperschneider, J., Nemri, A., Xu, B., Mago, R., Cuomo, C. A., Rathjen, J. P., Park,R. F., Ellis, J. G., and Dodds, P. N. (2015). Comparative genomics of Australian isolates of the wheat stem rust pathogen Puccinia graminis f. sp. tritici reveals extensive polymorphism in candidate effector genes. Frontiers in Plant Science, 5(January):759.

Wang et al., 2014. Wang, D., Ning, K., Li, J., Hu, J., Han, D., Wang, H., Zeng, X., Jing, X., Zhou, Q., Su, X., Chang, X., Wang, A., Wang, W., Jia, J., Wei, L., Xin,Y., Qiao, Y., Huang, R., Chen, J., Han, B., Yoon, K., Hill, R. T., Zohar, Y., Chen,F., Hu, Q., and Xu, J. (2014). Nannochloropsis genomes reveal evolution of microalgal oleaginous traits. PLoS genetics, 10(1):e1004094.

Weigel and Mott, 2009. Weigel, D. and Mott, R. (2009). The 1001 genomes project for Arabidopsis thaliana. Genome biology, 10(5):107.

with contributions from Andrew J. Bass et al., 2015. with contributions from Andrew J. Bass, J. D. S., Dabney, A., and Robinson, D. (2015). qvalue: Q-value estimation for false discovery rate control. R package version 2.4.2.

Zhao et al., 2012. Zhao, Y., Wu, J., Yang, J., Sun, S., Xiao, J., and Yu, J. (2012). PGAP: pan-genomes analysis pipeline. Bioinformatics (Oxford, England), 28(3):416–8.

